# Microbial metabolic specificity controls pelagic lipid export efficiency

**DOI:** 10.1101/2023.12.08.570822

**Authors:** Lars Behrendt, Uria Alcolombri, Jonathan E. Hunter, Steven Smriga, Tracy Mincer, Daniel P. Lowenstein, Yutaka Yawata, François J. Peaudecerf, Vicente I. Fernandez, Helen F. Fredricks, Henrik Almblad, Joe J. Harrison, Roman Stocker, Benjamin A. S. Van Mooy

## Abstract

Lipids comprise more than 20% of sinking organic matter in the ocean and play a crucial role in the carbon cycle. Despite this, our understanding of the processes that control lipid degradation is limited. Here we combined nano-lipidomics and imaging to study the bacterial degradation of diverse algal lipid droplets. Bacteria isolated from natural marine particles exhibited distinct dietary preferences, ranging from selective to promiscuous degraders. Dietary preference was associated with a distinct set of lipid degradation genes rather than with taxonomic origin. The top degrader, *Pseudomonas zhaodongensis,* rapidly consumed triacylglycerols (TAGs) from lipid extracts while promoting colonization of kin by chemotaxis toward glycerol, the TAG degradation product. Using synthetic communities composed of isolates with distinct dietary preferences, we demonstrated that lipid degradation is modulated by microbial interactions. A particle export model incorporating these dynamics suggests that metabolic specialization and community dynamics influences lipid transport efficiency in the ocean’s mesopelagic zone.

## Introduction

Carbon export in the ocean is mediated primarily by the sinking of particles composed of biomolecules and biogenic minerals originating from plankton. As these particles sink, they transport particulate organic carbon (POC), originally assimilated in the euphotic zone by phytoplankton through photosynthesis, from the surface to the deep ocean, a process known as the biological pump. The microscale interactions of heterotrophic organisms such as bacteria and zooplankton with sinking particles are critical mechanisms constraining the magnitude of the biological pump ^1^, for example, by transforming POC into dissolved forms within the upper water layer and thereby reducing the flux of carbon into the deep ocean ^2–4^.

Approximately 20% of surface ocean POC is composed of lipids ^5,6^, carbon-rich biomolecules used for energy storage, membrane structure, electron transport, and signaling ^7^. As particles sink from the euphotic zone to the deep sea, they are degraded by diverse communities of resident microbes ^8,9^, making particles hotspots of cell abundance and activity of hydrolytic enzymes, including lipases involved in lipid degradation ^10–12^. The flux of lipids decreases by an order of magnitude as particles descend through the mesopelagic zone ^5,13^ contributing to the microbial release of CO_2_ during remineralization. Thus, microbial lipid degradation on sinking particles exerts an important control on global CO_2_ concentrations. Understanding the microscale processes that mediate microbial lipid remineralization in the ocean is therefore important to improve our ability to forecast global carbon fluxes in changing ocean regimes.

Aside from their contribution to POC, lipids are powerful markers for understanding sinking particles in the ocean. For decades lipids were the primary biomarkers for assessing POC sources in the mesopelagic zone ^14,15^ and remain an important tool for understanding perturbations to the biological carbon pump ^16^. Many classes of lipids in plankton are highly labile and subject to molecular transformations that rapidly differentiate the lipid composition of sinking particles from that of living planktonic biomass ^17–19^. However, the mechanisms by which microbial communities degrade and ultimately remineralize lipids, and thereby attenuate the flux of this large component of POC through the mesopelagic zone, is poorly understood. Advanced marine lipidomics methods ^20,21^ have yet to be brought to bear on this important problem.

Here we combine nano-scale lipidomics with novel high-throughput fluorescence microscopy screening assays to show that bacteria isolated from natural sinking particles have distinct preferences for different classes of lipids. Using experiments with lipid droplets – models for lipid energy-storage bodies in phytoplankton cells – we demonstrate that different bacteria degrade different lipids at considerably different rates and, furthermore, that bacteria-bacteria interactions affect lipid remineralization rates. Using a mathematical model of sinking marine particles that incorporates bacterial degradation rates of lipid droplets, we show how microbial community structure could influence the transfer efficiencies of lipids from the euphotic zone to the deep-sea.

## Results and Discussion

### Lipid degradation dynamics are isolate-specific

The lipid droplets were composed of lipids extracted from a 400 L culture of nitrogen-starved *Phaeodactylum tricornutum* (CCMP632), a marine diatom that forms conspicuous amounts of storage lipids upon nitrogen starvation (Fig. 1A,B) ^22^. As marine plankton are typically under nitrogen limitation in the ocean ^23^, the composition of this lipid extract serves as a model for sinking aggregates formed by a declining diatom community. Using a recently developed, ultra-sensitive lipidomic pipeline ^21^, we found that the diatom lipid extract was dominated by triacylglycerols (TAGs), a class of lipid molecules that similarly dominates euphotic zone planktonic communities and sinking particles in the ocean ^13,24^. The extract also contained other ubiquitous lipid classes, including free fatty acids (FFAs), phospholipids (phosphatidylcholines, (PCs)), glycolipids (monogalactosyldiacylglycerols (MGDGs), digalactosyldiacylglycerols (DGDGs), sulfoquinovosyl diacylglycerols (SQDGs)) and chloropigments (chlorophyll *a* plus pheophytin *a*) (Fig. 1C).

**Fig. 1.**
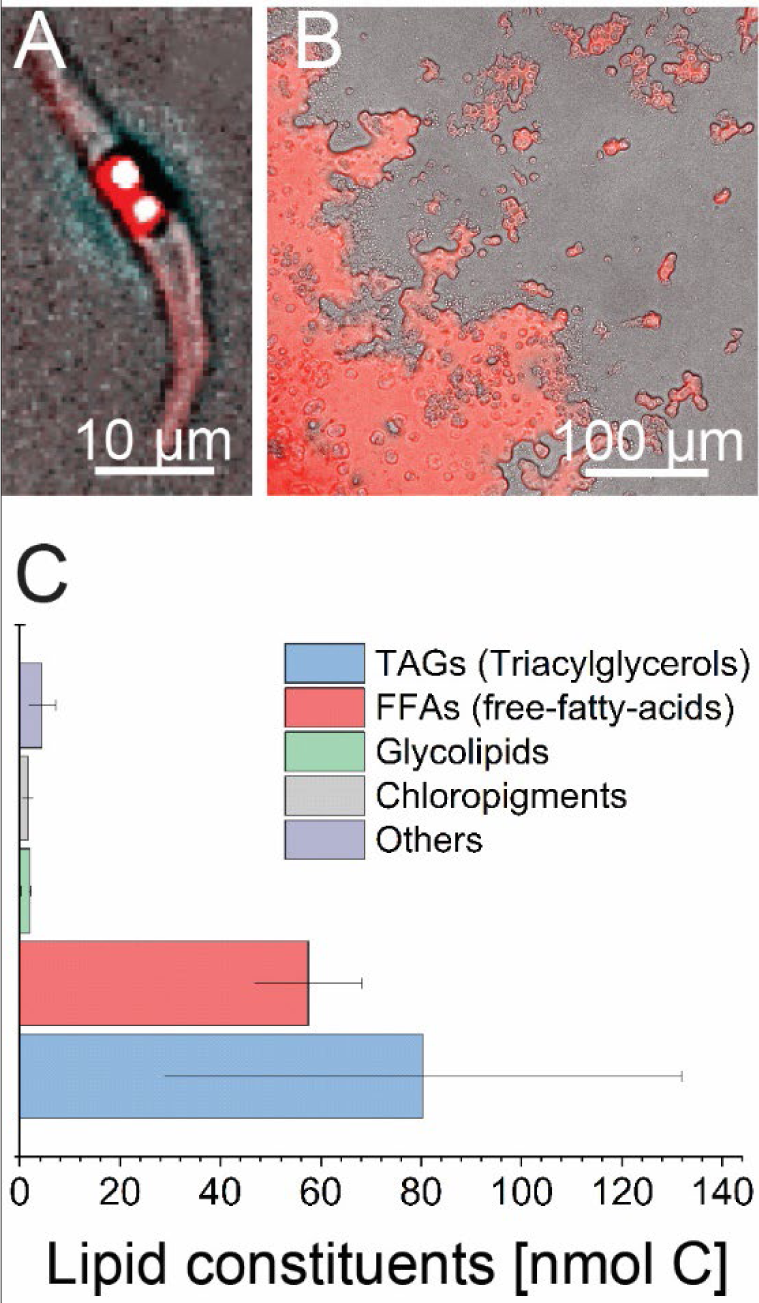
The lipid content of nitrogen-starved diatoms is dominated by triacylglycerols and free-fatty-acids. (**A**) A single cell of the marine diatom *Phaeodactylum tricornutum* stained with the lipid-specific stain Nile-red. (**B**) Lipid extracts from N-starved *P. tricornutum* cells adhere as droplets to a glass slide. Red fluorescence is due to the co-extraction of autofluorescent chloropigments. (**C**) The major lipid constituents of the extract from N-starved *P. tricornutum* cells. This lipid extract was used in all experiments unless indicated otherwise. Shown are the means (± SD) from five technical replicates.

We incubated individual droplets of diatom lipid extract (equivalent diameter 1.85 mm ± 0.21 mm, mean ± SD) with each of 24 isolates of bacteria: 21 isolates were grown from marine particles during a research cruise in Clayoquot sound (British Columbia) ^25^ and three are well-studied model isolates (*Ruegeria pomeroyi* DSS-3 (Rp3)*, Marinobacter adhaerens* HP15 (Ma1), and *Alteromonas macleodii* ATCC 27126 (Am2), see Table S1 for strain information). Lipid content in droplets was measured after 11 days of incubation (Materials and Methods), revealing that bacteria do not degrade lipids indiscriminately, but rather that specific groups of bacteria selectively consume certain lipids (Fig. 2). Four ‘dietary’ clusters emerged from this lipidomic analysis: isolates in cluster I (*n* = 6) promiscuously degraded all lipids; isolates in cluster II (*n* = 4) degraded all lipids except TAGs; isolates in cluster III (*n* = 9) degraded only FFAs, TAGs, and MGDGs; and isolates in cluster IV (*n* = 5) degraded only FFAs. Membership of the different dietary clusters did not correlate with the taxonomic origin of the isolates (Fig. S1). Instead, comparative genomics among four selected isolates from each dietary cluster (*A. macleodii* (Am2) from cluster I, *R. pomeroyi* (Rp3) from cluster II, *Pseudomonas zhaodongensis* (Pz15) from cluster III, and *Pseudoalteromonas shioyasakiensis* (Psh20) from cluster IV) revealed a distinct repertoire of putative TAG and fatty acid degradation (*fad*) genes (Fig. S2). Especially the copy numbers of *fad* gene paralogs, predicted to encode enzymes that catalyze the β-oxidation of prominent fatty acids (including free fatty acids; Fig. S2), varied greatly among the genomes of the sequenced isolates from the four dietary clusters. Together, these data indicate that marine bacteria have distinct dietary preferences for specific fractions of diatom lipids and that similar degrader phenotypes emerge from different genomic backgrounds rather than by taxonomic association.

**Fig. 2.**
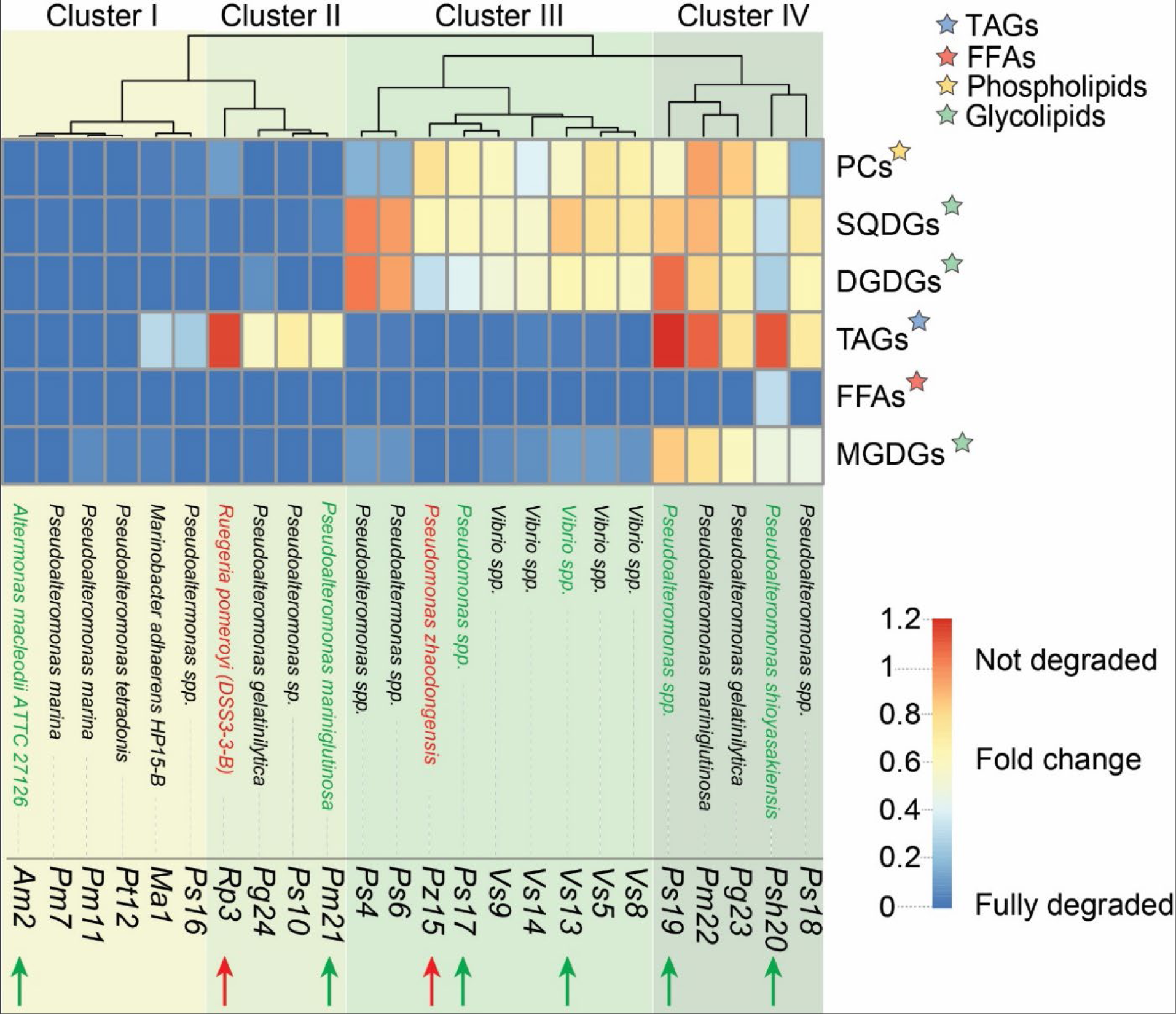
Marine bacteria exhibit distinct dietary preferences for different groups of phytoplankton lipids. Heatmap showing the degradation of different lipid fractions after incubation with bacterial monocultures (*n* = 24; species names and abbreviations indicated at the bottom). Shown are fold-changes in lipid content after 11 days of incubation. The different lipid constituents and their classification within major lipid groups are indicated by colored stars on the right of the heatmap. A nearest-neighbor clustering of the degradation profiles (i.e., the change in lipid content across the different lipid groups) revealed four clusters (I-IV), each characterized by different lipid fractions that were degraded to varying degrees by incubation with bacterial monocultures. Red arrows denote *Ruegeria pomeroyi* (Rp3) and *Pseudomonas zhaodongensis* (Pz15), two isolates selected for further biochemical and behavioral characterization (Figs. 3 and 4). Green arrows indicate additional isolates chosen as archetypical cluster representatives in experiments to quantify pairwise interactions when in co-culture with either *R. pomeroyi* (Rp3) or *P. zhaodongensis* (Pz15) (Fig. 5).

We next sought to determine whether bacterial dietary preferences affect the degradation kinetics of lipid droplets. We combined fluorescence microscopy with time-resolved lipidomic analysis (Videos S1-S3) and focused on the degradation behavior of two bacterial isolates distinguished by their divergent ability to degrade TAGs, the dominant lipid class in the extract from *P. tricornutum.* The two isolates were *R. pomeroyi* (Rp3), an isolate from cluster II which cannot degrade TAGs, and *P. zhaodongensis* (Pz15), an isolate from cluster III which degrades TAGs (Fig. 2, both isolates indicated by red arrows). Time-lapse microscopy imaging of lipid droplets incubated with *P. zhaodongensis* (Pz15) (Fig. 3A) revealed that the autofluorescence of chloropigments (chlorophyll plus pheophytin) in lipid droplets decreased by an average 19 ± 5.3 % (*n* = 5) within 15 h and remained low for the rest of the experiment. In comparison, in lipid droplets incubated with *R. pomeroyi* (Rp3) the autofluorescence decreased by only 4.4 ± 2.0 % after 42 h (*n* = 6), similar to the no-bacteria control (4.4 ± 1.0 %, *n* = 3, Fig. 3B).

**Fig. 3.**
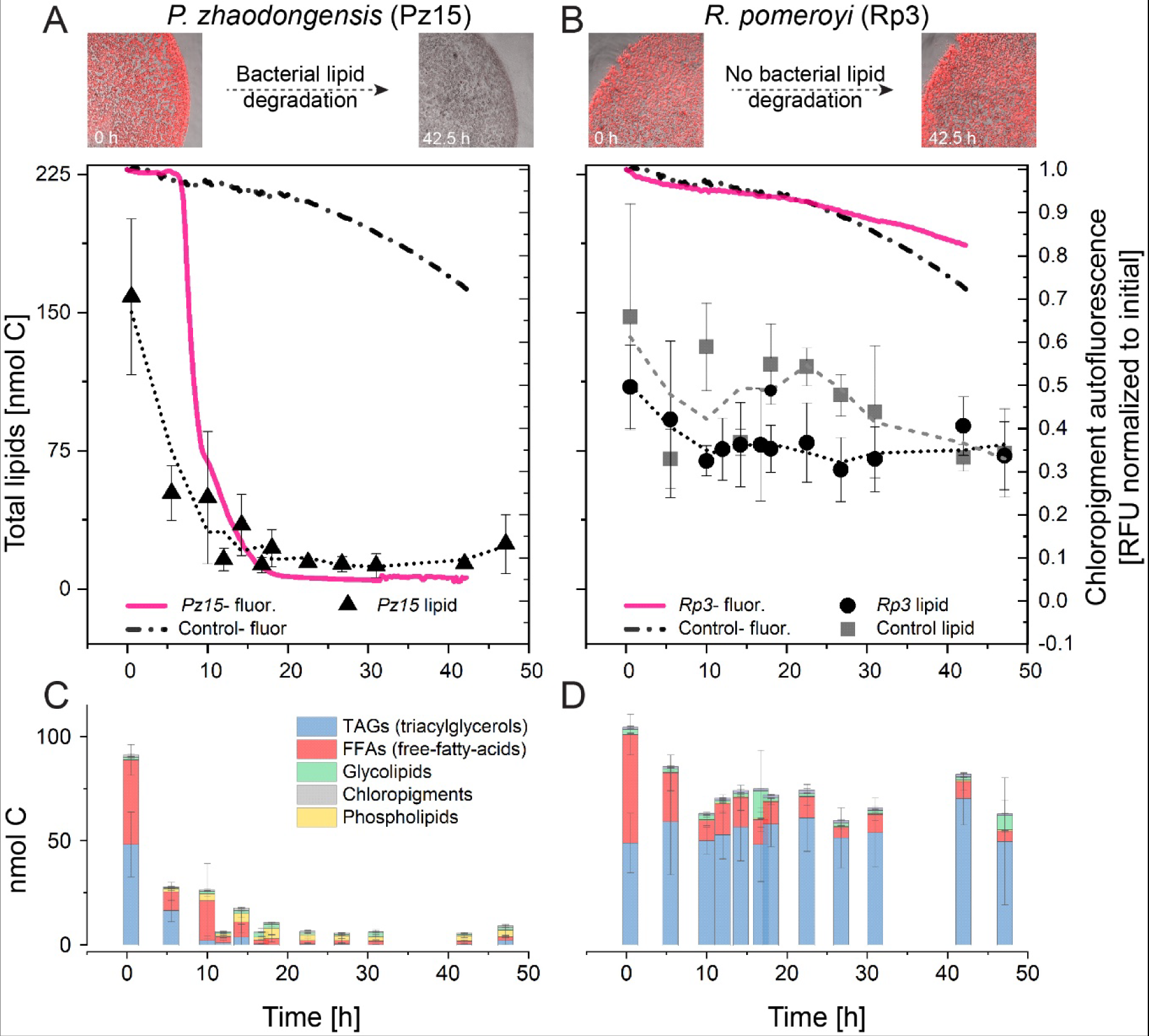
Marine bacteria have widely different lipid degradation kinetics. (**A, B**) The degradation of *P. tricornutum* lipid extracts over 48 h when incubated with *P. zhaodongensis (Pz15)* (left) or *R. pomeroyi* (Rp3) (right). Microscopy images show representative lipid droplets at the beginning of the time course (0 h) and near the end (42.5 h). For each time point, the total lipid content (left axis) and chloropigment autofluorescence (right axis) of individual lipid droplets was quantified using automated lipidomics and fluorescence microscopy, respectively. Average normalized fluorescence for bacterial treatments (pink solid line) and for background controls (black dashed line) are shown (*n* = 3, SD not displayed due to large variance). Total lipid content is shown as mean ± SD (*n* = 3) for *P. zhaodongensis* (Pz15, panel A, black triangles), *R. pomeroyi* (Rp3, panel B, black circles) and a no-bacteria control (panel B, gray squares). Lipid contents after microscopy image correction (caused due to sample handling, Fig. S5) are shown as smoothed dotted lines for *P. zhaodongensis* (Pz15, panel A, black dotted line), *R. pomeroyi* (Rp3, panel B, black dotted line) and a no-bacteria control (panel B, gray dotted line). (**C, D**) The lipidomic composition of lipid droplets in (A) and (B) upon incubation with *P. zhaodongensis* (Pz15, left) and *R. pomeroyi* (Rp3, right). Shown are the fractions of triacylglycerols (TAGs), free-fatty acids (FFAs), glycolipids, pheophytin and phospholipids.

Interestingly, despite the observed decrease in autofluorescence, the nano-lipidomics analysis revealed that chloropigments were not degraded by either isolate. We ascribe the loss of autofluorescence during the course of degradation to the well-documented concentration-dependent self-quenching of chlorophyll molecules in highly concentrated solutions (^26,27^, Fig. S3 and Video S4). Essentially, the relative concentration of chloropigments in the droplet increased as the TAGS, FAs, and other lipids were degraded, resulting in self-quenching of chloropigment autofluorescence that is correlated with the loss of lipids. Parallel analysis of lipid droplets using nano-lipidomics revealed that *P. zhaodongensis* (Pz15) degraded the total diatom-derived lipid pool almost completely within 48 h, from an initial 158 ± 42 nmol C (mean ± SD, *n* = 3 per time point) to 24 ± 16 nmol C, corresponding to an 84% reduction (Fig. 3A). By contrast, *R. pomeroyi* (Rp3) only degraded the total lipid pool from an initial 110 ± 22 nmol C to 72 ± 18 nmol C, corresponding to a 34% reduction (Fig. 3B). This difference in total lipid degradation between both isolates was also reflected in different amounts of phospholipids produced by the growing bacteria in their cell membranes ^28^. For *P. zhaodongensis* (Pz15), phospholipids increased from an initial 0.3 ± 0.1 nmol C to 5.4 ± 4.1 nmol C during 48 h, reflecting a 1490% increase (Fig. 3C), whereas for *R. pomeroyi* (Rp3) phospholipids increased from an initial 0.24 ± 0.1 nmol C to 0.7 ± 0.0 nmol C during 48 h, corresponding to a 220% increase (Fig. 3D). We note that *R. pomeroyi* (Rp3) also produces ornithine lipids, which were not rigorously quantified as part of this study. Combined chloropigment autofluorescence and lipidomic measurements (Fig. 3A, B) revealed that quenching of chloropigment autofluorescence is a good proxy for lipid degradation and that the total content of lipids in a droplet decreased linearly with decreasing fluorescence intensity (Figs. S4-S5, Supplementary Materials and Methods). Together, these data on both isolates demonstrate that differences in their dietary preferences for lipids have a major impact on rates of lipid droplet degradation and, consequentially, on the isolates’ growth rates.

In parallel with time-resolved fluorescence microscopy, our nano-lipidomic analysis enabled us to track the sequence of TAG and FFA degradation by *R. pomeroyi* (Rp3) and *P. zhaodongensis* (Pz15) within single droplets of lipid extract (Fig. 3C,D). Here *P. zhaodongensis* (Pz15) degraded TAGs from an initial 80 ± 25 nmol C to 0.8 ± 0.7 nmol C during the first 16 h (a 99% reduction), while concurrently also degrading FFAs (initial 67 ± 12 nmol C to 3.4 ± 0.6 nmol C, a 94% reduction; Fig. 3C). In contrast, during the first 16 h, *R. pomeroyi* (Rp3) did not degrade TAGs (49 ± 14 nmol C to 48 ± 18 nmol C), but degraded FFAs (initial 52 ± 10 nmol C to 12 ± 4 nmol C, a 77% reduction; Fig. 3D). These results recapitulate the findings from our initial screening of isolates (Fig. 2), and additionally show that *P. zhaodongensis* (Pz15) expresses its specific lipid degradation enzymes even at the initial stages of degradation. Thus, the dietary preference for different classes of lipids appears to be dictated by the inherent metabolic capabilities of each isolate and, at least in this study, not by the changing composition of the lipid droplet upon degradation. To better understand the origin of the dietary preference for these two isolates, we constructed and compared lipid degradation pathways *in silico* from the genomes of *P. zhaodongensis* (Pz15) and *R. pomeroyi* (Rp3) (Fig. S2). We found that the two isolates had distinct sets of lipid degradation enzymes, including a different set of acyl-CoA dehydrogenases. In addition, *P. zhaodongensis* (Pz15) possesses a triacylglycerol lipase, which is necessary for TAG degradation, whereas *R pomeroyi* (Rp3) lacks this enzyme. By contrast, both *P. zhaodongensis* (Pz15) and *R. pomeroyi* (Rp3) genomes possess multiple paralogs for genes encoding functions that are predicted to drive FFA degradation.This suggests that the differences in lipid degradation are primarily associated with differences in the genetic repertoire of the two isolates (Fig. S2).

The onset of rapid lipid degradation by *P. zhaodongensis* (Pz15) often occurred after a time delay, typically of 7-12 h (Fig. 3A). To understand the origin of this delay, we engineered *P. zhaodongensis* (Pz15) to constitutively express GFP from a plasmid (Supplementary Materials and Methods) and used fluorescence microscopy to investigate the microscale dynamics of lipid droplet degradation. This investigation revealed that only a small number of *P. zhaodongensis* (Pz15) cells immediately attached to the droplet upon inoculation, whereas most of the colonization was delayed (Video S5). When GFP-labeled *P. zhaodongensis* (Pz15) cells were provided with a droplet composed of a synthetic TAG mixture (tripalmitoleic acid, three 16:1 fatty acids), motility and chemotaxis became prominent after a 10 h delay and motile bacteria formed high concentrations of bacterial clusters on the droplet (Video S6). We attribute this behavior to the degradation induced release of the glycerol moiety from TAGs to which *P. zhaodongensis* is chemoattracted (Fig. S6) and which is likely released during the cleavage of TAGs. This result highlights that early bacterial colonizers are important for lipid degradation not only because of direct degradation, but also because they can attract further degraders through the release of degradation products or metabolites that can act as chemoattractants ^29^.

### Microbial interactions modulate the rate of lipid remineralization

We next set out to examine whether the correlation between dietary preference and degradation rate in *P. zhaodongensis* (Pz15) and *R. pomeroyi* (Rp3) extended to other bacteria isolated from particles. To do so we chose six additional isolates as representatives of the different dietary clusters (Fig. 2, green arrows) and measured their degradation dynamics in shorter-term experiments (48 h). The relationship between chloropigment autofluorescence and total lipids allowed us to use fluorescence as a time-resolved readout of lipid degradation in a high-throughput fluorescence plate-reader assay (Figs. S4-S5, Supplementary Materials and Methods). From the resulting fluorescence curves we could estimate the delay before the onset of degradation and the maximum degradation rate (Fig. S7). By plotting the maximum degradation rate against the delay in the onset of degradation (Fig. 4), we found that the selected isolates largely clustered according to their dietary preference previously determined in lipidomic experiments (Fig. 2). The fastest degradation rate (7.0 ± 0.9 nmol C h^-1^, *n* = 10) was exhibited by *Alteromonas macleodii* (Am2, Fig. 4), belonging to dietary cluster I (i.e., degrading all lipid constituents; Fig. 2). Slower degradation rates in the range 2.3–5.3 nmol C h^-1^ (mean 3.8 ± 0.9 nmol C h^-1^, *n* = 26) were exhibited by *P. zhaodongensis* (Pz15), *Vibrio spp*. (Vs13) and *Pseudomonas spp*. (Pf17) (Fig. 4), belonging to dietary cluster III (i.e., degrading three out of the six detectable lipid classes: FFA, TAGs and MGDGs; Fig. 2). These three isolates showed delay times in the range 7.8–17.3 h (13.0 ± 2.6 h, *n* = 26). The longest delay (>48 h) was exhibited by two *Pseudoalteromonas* isolates (Ps19, Psh20), belonging to dietary cluster IV (i.e., only degrading FFAs; Fig. 2). We note that this fluorescence plate-reader assay could not resolve the short-term lipid degradation kinetics of the most fastidious isolates. Specifically, we observed lipid degradation by *R. pomeroyi* in only 3 out of 18 replicates (Fig. 4), while no measurable degradation occurred for *P. mariniglutinosa* (Pm21) over the 48 h experiment. This suggests that *R. pomeroyi* (Rp3) and *P. mariniglutinosa* (Pm21) require longer timescales to degrade lipids fully and are not as capable of degrading lipids over shorter timescales, a result that, for *R. pomeroyi* (Rp3), is consistent with our earlier experimental evidence (Fig. 3B,D). Overall, the recapitulation of the dietary clusters in bacterial degradation kinetics suggests that lipid degradation by single isolates in the marine environment is fundamentally linked to their dietary preference.

**Fig. 4:**
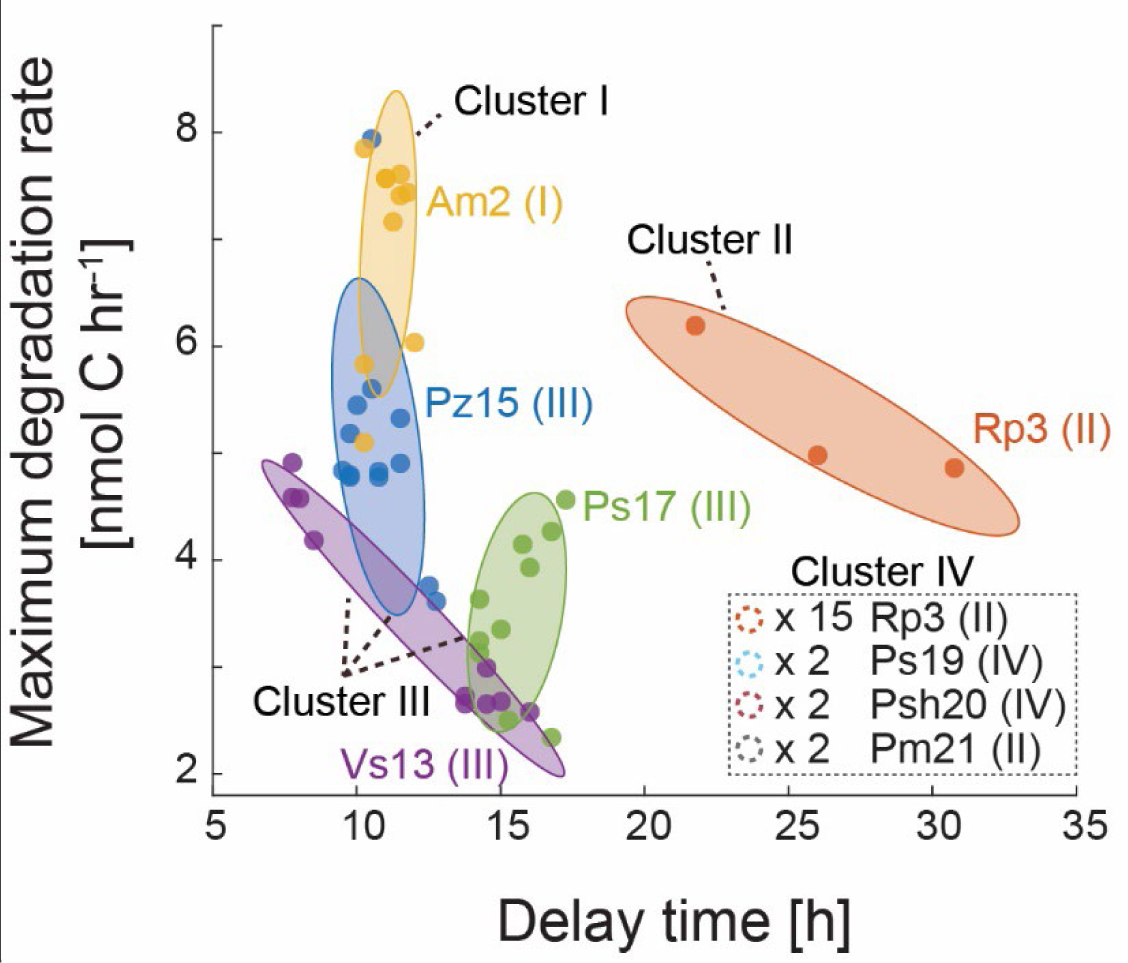
The lipid degradation kinetics of single bacterial isolates recapitulate their long-term degradation behavior. The kinetics of lipid droplet degradation derived from fluorescence time series for eight bacterial isolates (Pz15, Rp3, Vs13, Ps17, Am2, Ps19, Psh20 and Pm21) in monocultures, taken to be representative of the different dietary clusters (see Fig. 2, green arrows). Cluster affiliation is indicated following the descriptor of the bacterial isolate (e.g., ‘Pz15 (III)’, indicates that *P. zhaodongensis* placement into cluster III). Maximum degradation rates and delay times were measured in two independent experiments (*n* = 4–10 for each isolate, designated by individual dots). Interrupted circles in boxed legend (Cluster IV, bottom right) denote isolates with delay times greater than 48 h. Ellipsoids indicate the members of the dietary clusters from Fig. 2 and are labeled according to their cluster affiliation.

To understand how dietary preference also affects degradation rates by multispecies communities of bacteria, which are ecologically more relevant than species in isolation, we investigated the effect of pairwise interactions on lipid droplet degradation via the plate-reader assay (Supplementary Material and Methods). In experiments in which either the fast-degrader isolate *P. zhaodongensis* (Pz15) or slow-degrader *R. pomeroyi* (Rp3) were inoculated together with single representative isolates chosen from the four dietary clusters (Fig. 2, green arrows), we found that simple synthetic co-cultures exhibited different degradation rates and delay times when compared to monocultures at the same cell density. Synergistically faster lipid degradation rates (student’s *t*-test, *p* < 0.05) were observed when *P. zhaodongensis* (Pz15) was combined with *A. macleodii* (Am2), *Vibrio spp*. (Vs13) or *Pseudomonas spp*. (Pf17) (Fig. 5A, *n* = 6-7 per combination), resulting in a 16–57% higher maximal degradation rate when compared to normalized monoculture rates (Materials and Methods). In contrast, no significant changes in maximum degradation rates were observed in pairwise interactions that included *R. pomeroyi* (Rp3) (Fig. 5B, *n* = 9-10 per combination). The time delay was also affected by pairwise interactions. We found significantly shorter delays when the isolates were co-cultured with *P. zhaodongensis* (Pz15) when compared to their respective monocultures (Fig. 5C, *n* = 6-7 per combination). In contrast, *R. pomeroyi* (Rp3) co-cultures showed consistently longer delay times (Fig. 5D, *n* = 9–10 per combination).

**Fig. 5:**
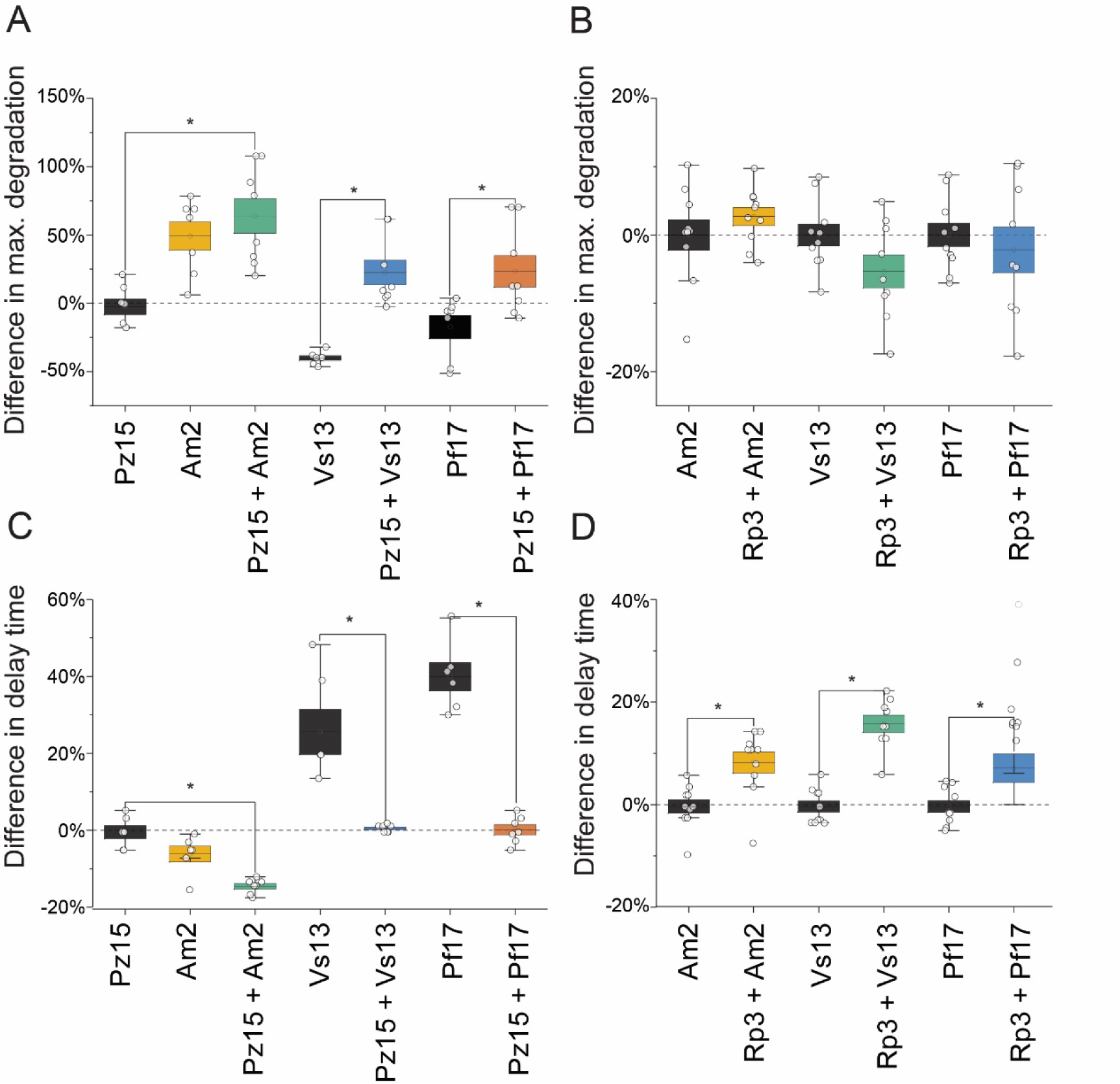
Pairwise interactions among marine bacteria affect the maximum rate and delay time of lipid degradation. The kinetics of lipid droplet degradation for synthetic two-isolate communities containing *P. zhaodongensis* (Pz15) or *R. pomeroyi* (Rp3) as focal isolates. (**A**) Two-isolate communities containing *P. zhaodongensis* (Pz15) demonstrate an increase in maximum degradation rates when compared to a monocultures of *P. zhaodongensis* (Pz15)*, Vibrio spp.* (Vs13) or *Pseudomonas spp*. (Pf17). The maximum degradation rate is calculated as the slope of a linear regression line fitted between the onset of degradation and its completion (Materials and Methods, Fig. S7). All rates are normalized to the maximum degradation rate in monoculture experiments and expressed as percent change. (**B**) Two-isolate communities containing *R. pomeroyi* (Rp3) show no significant difference between maximum degradation rates when compared to the respective monocultures of *A. macleodii* (Am2), *Vibrio spp.* (Vs13) or *Pseudomonas spp*. (Pf17). (**C**) Two-isolate communities containing *P. zhaodongensis* (Pz15) show a reduction in delay time when compared to a monoculture of *P. zhaodongensis* (Pz15)*, Vibrio spp.* (Vs13) or *Pseudomonas spp*. (Pf17). The delay time was defined as the time from the start of the experiment to the onset of degradation (Fig. S7). (**D**) Two-isolate communities containing *R. pomeroyi* (Rp3) show an increase in delay time when compared to the respective monocultures of *A. macleodii* (Am2), *Vibrio spp.* (Vs13) or *Pseudomonas spp*. (Pf17). In all graphs, the white dots represent individual measurements (*n* = 6–10 per condition). An asterisk indicates statistically significant differences (student’s t-test*, p* < 0.05) between treatments.

The results from these experiments indicate that specific members of simple communities can accelerate or delay the overall lipid degradation compared to single isolates. To gain further insight into the nature of these interactions, we used fluorescence microscopy to investigate lipid degradation by GFP-labeled *P. zhaodongensis* (Pz15) co-cultured with *A. macleodii* (Am2), a combination that yielded the shortest delay time and highest maximum degradation rate among all combinations of isolates tested (Fig. 5). For this simple consortium, we observed a well-mixed distribution of both isolates on the droplet (Video S7), potentially indicating syntrophy and the sharing of public goods ^30^ rather than e.g., the formation of segregated patches. *Pseudomonas* relatives are known to be nutritionally versatile and to metabolize many carbon-rich compounds, including crude oil, oil derivatives ^31^ and aliphatic compounds ^32^, which suggests that simultaneous exploitation of similar lipid constituents-rather than a sequential consumption of each constituent by itself-could have caused faster degradation rates and shorter delay times in communities that include *P. zhaodongensis* (Pz15). Pairwise interactions with Rp3 (Fig. 5D) led to extensive delays in degradation. These delays are similar to those observed in interactions between secondary and primary degraders of simple synthetic polysaccharide particles ^33^, and may be due to the physical blockage of the lipid surface or, alternatively, due to the production of secondary metabolites – a common phenomenon in Rhodobacter species ^34,35^. Our results, based on complex mixtures of compounds from plankton-derived lipid extracts, demonstrate that dietary preferences and microbial interactions at the scale of individual particles together control the rate of lipid remineralization by marine bacteria.

### Potential implications of microbial interactions on oceanic lipid transport

Using our experimentally measured lipid droplet degradation dynamics, we developed a mathematical model to assess the potential effect of microbial interactions on vertical lipid transport in the ocean (Supplementary Model, Fig. S8). The model tracks the sinking of individual particles from a size distribution that reflects the marine environment. Each particle contains a fixed proportion of lipid droplet and ballast and sinks at a speed that depends on its size and density as it descends from the base of the euphotic zone. Our model differs from recent single-particle approaches ^3,36^ in that it considers two distinct phases within a particle: an lipid droplet phase (density = 8.8 × 10^5^ g m^-3^) that can be degraded by bacteria and a ballast phase (density = 2.35 × 10^6^ g m^-3^) that cannot. As lipid, but not ballast, is consumed degradation leads to an increase in the particle density and thus sinking speed. As lipids are immiscible with water, likely limiting bacterial growth and degradation to the lipid–water interface ^37–39^, we assume a pseudo-first-order rate law, where the absolute rate of lipid degradation is dependent on the surface area of the lipid droplet phase exposed to colonization. To calculate a range of possible values for the lipid degradation rate per unit of exposed lipid droplet surface (*k_A_*_;_ mol C m^-2^ s^-1^), we combined the observed total degradation rates from our experiments with multiple isolates (Fig. 4; expressed as mol s^-1^ and dependent of the size of the observed droplet) together with our data of lipid droplet surface area (m^2^) from the same experiments. Since these two datasets are not paired, i.e. degradation rate and surface area were not measured on the same droplet, we used a Monte-Carlo approach to sample both datasets in order to obtain a representative distribution of degradation rates per unit area *k_A_* (Fig. S8A). We run the model using a range of bacterial degradation rate constants (*k_A_* = 2 × 10^-6^ to 1 × 10^-4^ g m^-2^ s^-1^) and a range of degradation delay times (*t*_delay_ = 0, 6, 12, 24 h), spanning the ranges observed in our experiments for these parameters. By simultaneously tracking the degradation of the lipid phase in each particle and the depth of the particle (as determined by its sinking speed), the model computes the particle concentration and the lipid content per particle at any given depth. From this, we quantify the transfer efficiency of the vertical lipid flux, defined as the ratio of the lipid flux at depth *z* relative to the initial flux at *z_0_*, for each class of particle size and over the entire particle population. The lower the transfer efficiency, the stronger is the attenuation of the lipid flux due to degradation by bacteria of the lipid phase.

The model predicts how microbial lipid degradation rates and delay times affect the lipid transfer efficiency of particles of different diameters containing lipid droplets. For smaller particles, the transfer efficiency is controlled almost exclusively by the degradation rate. Even when lipid degradation proceeds at the low end of the range of observed rates (*k_A_* = 2 × 10^-6^ g m^-2^ s^-1^), less than half of the lipid flux carried by particles smaller than 100 um diameter survive the descent to the bottom of the mesopelagic (1,000 m depth). Furthermore, the delay in the onset of degradation has almost no effect on the transfer efficiency of small particles containing lipid, which can be interpreted as a direct consequence of their very slow sinking speed: for particles of 100 µm diameter having an initial sinking speed of ∼12 m day^-1^, even the largest delay (= 24 h) remains small compared to the time it takes for the particles to reach depths of hundreds of meters, and so the amount of degradation with and without delay is similar. By contrast, the delay has a much greater effect on rapidly degrading, larger particles which rapidly sink to depth. For example, if degradation is delayed by 24 h, a particle with a diameter of 250 μm sinks almost unscathed through the top 100 m (due to its initial sinking speed of ∼80 m day^-1^) even when the highest degradation rate is considered. Conversely, without any delay, 20% of the lipid in the particle will degrade over the same sinking distance of 100 m. Our model also provides estimates of total lipid transfer efficiencies (TE) at a depth of 1,000 m (TE_900_; *z_0_* = 100 m; *z* = 1,000m) for the total lipid flux summed across initial droplet sizes; we found the TE_900_ ranges from ∼97% for the lowest degradation rate and longest delay to TE_900_∼57% TE at the highest rate and no delay. This strong variation of transfer efficiency (more than 10x more attenuation at the highest degradation rate than at the lowest) results from the wide range of degradation rates considered, itself stemming from varying synergistic or antagonistic impacts of the community’s composition on degradation. While our mathematical model does not provide a comprehensive description of all factors involved in lipid flux attenuation (e.g., particle disaggregation) and is based on rates obtained *in vitro* rather than from the environment, it illustrates the importance of particle colonizers in determining the fate of lipids within sinking particles. The model also provides mechanistic insight into which classes of particle size will be more sensitive to variations in degradation rates and the delay of degradation. For example, given that the observed transfer efficiency of both TAGs and free fatty acids through the mesopelagic is less than 10% in field studies ^5,13^, our model evokes the hypotheses that synergistic bacterial lipid degradation is common and that particles containing lipid droplets in the marine environment are more likely to be small and sink slowly.

By coupling the quantitative analysis of physical processes to the effects of bacterial community structure on chemical processes, on the scale of individual particles, our study provides new insights into the degradation of sinking POC. Furthermore, the approach taken here – elucidating the connections between the molecular composition of POC, dietary preferences of bacteria for different molecules, and degradation kinetics – provides a blueprint for the study of the fate of phytodetritus, fecal pellets, and other organic carbon entities in the ocean. Results from such well-controlled, laboratory-based studies provide the optimal ingredients to achieve a more mechanistic understanding of particle degradation and thus derive fundamental principles and laws that can inform large-scale models of the ocean carbon cycle.

## Materials and Methods

### Phytoplankton growth

Cultures of *Phaeodactylum tricornutum* CCMP2561 were grown in a volume of 400 L of seawater based modified L1+Si medium in the facilities of the National Center for Marine Algae and Microbiota (NCMA, Bigelow, ME, USA). To generate N-starved cultures, L1 media was prepared as per standard recipe but NO_3_ concentrations were adjusted to 1/16th of normal and Si concentrations to 2 X normal. Small inoculum volumes were used to avoid media carryover which could potentially increase final NO_3_ concentrations. Upon reaching mid-exponential growth, cells were harvested by continuous centrifugation and the resulting pellet was frozen at -80°C, shipped on dry-ice and used as input material in lipid extraction protocols (below).

### Bacterial isolation, growth media and physiological buffers

All marine bacterial isolates and strains are listed in Table S1. Marine aggregates were collected in particle traps during a scientific research cruise in Clayoquot sound (British Columbia, Canada) ^25^. Collected marine aggregates were spread onto tryptone seawater agar plates (2% agar, 0.1% tryptone) and the resulting colonies were streaked to axenity on similar plates. Individual clones were grown at 30°C in marine broth 2216 (Sigma Aldrich, 76448) augmented with sea salt to a salinity of 36 psu (Instant Ocean, Blacksburg, VA, USA) and agitated at 200 rpm before cryopreservation at - 80°C with 10% DMSO. For experiments, cryopreserved bacteria were grown overnight in marine broth 2216 at 30 °C and 200 rpm agitation. The resulting culture was centrifuged at 2350 g for 2 min and washed with sterile seawater based f/2 medium (36 psu). This process was repeated 3 times and the washed culture was incubated at 30 °C for a minimum of 2 h. After this starvation period, the optical density at 600 nm was measured on a spectrophotometer (BioTeK, Synergy HTX Multi-Mode Reader) and adjusted to 0.01 (corresponding to ∼1 × 10^7^ mL^-1^). The final inoculation concentration of all bacterial cells in experimental setups was obtained by diluting the above washed culture into fresh f/2 medium. The concentrations of cells was either 2 × 10^6^ cells mL^-1^ for the pairwise interaction experiments (see *Pairwise interaction experiments*) or 1 × 10^6^ cells mL^-1^ for all other experiments. Where appropriate, bacteria were grown and propagated on lysogeny broth (LB), which contained, per liter: 10 g tryptone (Difco), 5 g yeast extract (Thermofisher Scientific), 5 g NaCl (Thermofisher Scientific). Semi-solid agar media was prepared by adding 15 g L^-1^ bacto agar (Difco, Fisher Scientific) to LB or marine broth 2216.

Alternatively, Zobell Marine Agar 2216 was purchased (Himedia, Mumbai, India) and prepared according to manufacturer’s directions. Phosphate buffered saline (PBS) was purchased as a 10 X concentrated solution (Amresco, Solon, OH, USA) and diluted, as required, in sterile MilliQ-treated water.

### Lipid extraction and droplet spotting

Bulk lipids were extracted from frozen, pelleted cells of *Phaeodactylum tricornutum* via a Bligh & Dyer extraction protocol ^40^. The resulting phytoplankton lipid extract was highly viscous, and lipid extracts were diluted in 99.5% Dichloromethane (Sigma Aldrich, D65100) at a w/v ratio of 1:100 to facilitate the dispensation of small volumes. Dispensation was done using a 100 µl glass Syringe (Hamilton 710N series, Hamilton Corp., Reno, NV, USA) connected to a 10 µl PCR glass micropipette (Drummond, Broomall, PY, USA) via Tygon (ND-100-80, Saint-Gobain Performance Plastics, Paris, France) tubing filled with deionized water. A PicoPlus syringe pump (Harvard apparatus, Holliston, MA, USA) was used to pre-load the glass micropipette with 10 µl of diluted lipid extracts (at all times avoiding contact with plastic surfaces) and then used to spot lipid droplets by setting a 1 µl s^-1^ dispensation rate and a dispensation volume of 400 nl. Using this setup, lipid droplets were spotted by placing the tip of the glass micropipette in contact with the receiving glass surface and rapidly retracting the pipette. This resulted in lipid droplets with an equivalent diameter of 1.85 ± 0.21 mm (SD) and a height of ∼ 10 µm as determined by confocal microscopy (Zeiss confocal, LSM 510 META). Lipid droplets were either spotted into the bottom of low-absorption LC-MS vials (Supelco, Bellefonte, PA, USA) or glass bottom 96-well plates (Cellvis, Mountain View, CA, USA), depending on the experimental setup. Residual DCM was removed from the phytoplankton lipid by placing vials or plates under vacuum for >1h and lipid integrity was preserved by placing plates/vials with spots into -80 °C and using them within a day.

### Lipid droplet visualization

Lipid droplets were found to strongly autofluoresce under an excitation/emission combination typically used for Cy5 imaging (Ex.: 628/40 nm HBW, Em.: 692/40 nm HBW). This autofluorescence is caused by co-extracted photopigments, i.e. chlorophyll *a* and *c* ^41^, and their chemical conversion to pheophytin. We exploited the natural lipid autofluorescence to estimate the degradation of lipids (see section on *Lipid autofluorescence quenching as a proxy for lipid degradation*) and to visualize the bacterial interactions occurring on lipid droplets. All lipid droplets were visualized on an automated fluorescence microscope (Nikon-Eclipse Ti, Nikon Corp., Tokyo, Japan) with excitation wavelengths provided by a Spectra X light engine (Lumencor, OR, USA). Temperature control was provided by an encasing incubator (Life Imaging Services, Basel Switzerland) and kept constant at 30°C for all experiments.

### Initial screening of bacterial isolates for lipid degradation

In order to determine the ability of microbial isolates (*n* = 24, see *Bacterial isolation and cultivation*) to degrade phytoplankton lipids, isolates were incubated in the presence of lipid droplets. For this, cells were dispensed into LC-MS vials (to a concentration of 1×10^6^ ml^-1^) containing lipid droplets and hereafter incubated at 30 °C. Immediately after dispensation, a first set of vials were sampled and frozen at - 80 °C (= *C*_*_0_* time point) and the remainder incubated for a total duration of 11 days before final endpoint sampling (= *C*_*_11_* time point). All samples were kept at -80 °C after sampling and shipped in liquid N_2_ filled dewars for chemical processing.

### Combined microscopic imaging and lipidomic sampling

In order to combine physical observations with chemical sampling, LC-MS vials with lipid droplets and selected bacterial isolates were placed into a microscopy-compatible custom-built acrylic holder (Fig. S9). This permitted the simultaneous microscopic imaging of dispensed lipid droplets and bacteria at the bottom of the vials and their regular sampling for chemical fixation. Due to space constraints on, additional LC-MS vials were placed adjacent to the microscopy stage under identical temperature conditions and collected for lipidomic analysis at identical times.

### Lipidomic analysis

All lipidomic analysis was done according to a previously described method ^21^. In short, nanoflow high-performance liquid-chromatography (nano-HPLC) was performed with a Thermo Scientific Easy nLC 1200 apparatus. Samples were injected in a direct mode onto a Thermo Scientific Acclaim PepMap 100 (75 μm × 2 cm; 3 μm; 100 Å), C18 guard column, and a Thermo Scientific Acclaim PepMap RSLC (75 μm × 15 cm; 2 μm; 100 Å) C18 analytical column, housed in a Phoenix S&T PST-BPH-20 butterfly column heater operated at 50 °C. The samples were dissolved prior to analysis in 50% water and 50% isopropanol. Eluent A consisted of 75% water, 25% acetonitrile, 0.1% formic acid, and 0.04% ammonium hydroxide; Eluent B consisted of 75% isopropanol, 25% acetonitrile, 0.1% formic acid, and 0.04% ammonium hydroxide. The chromatographic gradient profile, at a constant flow rate of 300 nL min^-1^, is defined in ^23^. Thermo Scientific Nanospray Flex source and stainless-steel emitter tip were used to couple the nano-HPLC system to a Q-Exactive orbitrap mass spectrometer. Data processing and annotation were performed using the LOBSTAHS ^23,42^.

### Lipid autofluorescence quenching as a proxy for lipid degradation

The quantification of lipid autofluorescence in parallel with nano-LC-MS lipidomic analysis revealed that the total amount of lipid present was approximately proportional to the magnitude of fluorescence (Fig. S4). Lipid fluorescence quenching is mediated by the bacterial degradation of lipids and the concurrent accumulation of co-extracted pheophytin, i.e. chlorophyll without Mg^2+^ ion. As some bacteria in our collection degraded lipids, but not pheophytin, the accumulation of pheophytin leads to a loss of autofluorescence by concentration dependent quenching mechanisms ^26,43^. These relationships (between measured fluorescence and chlorophyll concentration and between lipid content and chlorophyll concentration) were parameterized directly from experiments (Fig. S4) and allowed us to relate both parameters and establish fluorescence as a non-invasive readout of lipid degradation.

To quantify lipid degradation, time series data was generated for Cy5 fluorescence data (Ex.: 628/40 nm HBW, Em.: 692/40 nm HBW) alongside abiotic droplet controls. Through comparison, the total bacterially degraded lipids were estimated (Fig. S7). A smoothing spline with a five hour window was then applied, identifying the time offset (delay) and magnitude of the highest degradation rate (Fig. S7). This technique was applied to fluorescence dynamics for individual isolates, as well as for multispecies communities to enable a rapid screening assay.

### Pairwise interaction experiments

In order to determine whether the interaction between bacterial isolates affects the degradation of phytoplankton lipid extracts, selected pairs of microbial isolates co-incubated in the presence of lipid droplets. For this, lipid droplets were spotted into glass-bottom 96-well plates (see *Lipid extraction and droplet spotting*) and the isolates *P. zhaodongensis* (Pz15) or *R. pomeroyi* (Rp3) were combined with selected bacterial isolates from the strain collection. Each isolate was added at a concentration of 1 × 10^6^ ml^-1^, always resulting in final concentrations of 2 × 10^6^ ml^-1^ per reaction volume. For single-isolates in interaction experiments, the amount added was doubled to achieve a final concentration of 2 × 10^6^ ml^-1^ per reaction volume. Using a fluorescence plate reader (BioTeK, Synergy HTX Multi-Mode Reader), the fluorescence decay of lipid droplets was observed using Cy5 specific excitation/emission (Ex.: 620/40 nm HBW, Em.: 680/30 nm HBW). Fluorescence was assessed at the bottom of each well every 15 minutes for a total duration of 48 h.

### Chemotaxis experiments

Chemotaxis of *P. zhaodongensis* (Pz15) towards glycerol was measured using a modified capillary assay ^44^. Cultures of Pz15::GFP (see *Electroporation and GFP expression*, below) were incubated overnight in marine broth 2216 augmented with 100 nM of glycerol and then washed and processed as previously described ^44^. Glass capillaries were heat-sealed on one end and, before cooling, introduced into a sterile f/2 solution augmented with 10 µM glycerol. To observe the chemotactic behavior under the microscope, scotch tape was wrapped around a microscopy slide eight times to obtain a thickness of ∼1 mm and thereafter cut using a scalpel to create a rectangular ‘pool’ for bacterial visualization. The loaded capillary was placed so it entered the rectangular pool and was fixated with scotch tape. After preparation, the resulting slide was placed onto an inverted fluorescence microscope and 100 µl of *P. zhaodongensis* (Pz15) culture was carefully pipetted into the pool. After 15 min incubation period, swimming behavior around the capillary was recorded by capturing 500 frames at a frame rate of 10 per second. The resulting maximum intensity images captured the motility tracks for individual cells. To observe background motility of *P. zhaodongensis* (Pz15), maximum intensity images were created as far away as possible from the capillary (3 mm).

### Standard molecular methods

All basic microbiological and molecular procedures were executed according to standard protocols ^45^. Oligonucleotide primers were purchased from Integrated DNA Technologies. Plasmid preparations and PCR purifications were performed using kits purchased from Qiagen or BioBasics. Taq DNA polymerase and RNase A were purchased from New England Biolabs. Phusion DNA polymerase was purchased from ThermoFisher Scientific. Sanger sequencing was provided by the University Core DNA Services at the University of Calgary (https://www.ucalgary.ca/dnalab/).

### Identification of marine bacterial isolates by 16s rRNA gene sequencing

Bacterial 16s rRNA genes were amplified using a colony PCR protocol with 27F (5ʹ-AGAGTTTGATCCTGGCTCAG-3ʹ) and 926R (5ʹ-CCGTCAATTCMTTTRAGT-3ʹ) primers, which target the variable V1, V2 and V3 regions of the 16s rRNA gene. PCR products were purified from reaction mixtures and sent for Sanger sequencing using the 27F and 926R primers. Forward and reverse reads were manually trimmed for quality (i.e. only unambiguous base-calls from chromatograms were included). The bidirectional reads were then assembled using the Geneious Prime (2020) De Novo Assembly algorithm with default settings (highest sensitivity) ^46^. The assembled sequences were then queried against the curated EZBioCloud database of 16s rRNA gene sequences ^47^ (version 20200225, accessed March 19, 2020). We made presumptive genus-level identification based on a query sequence that had either (i) >98.7% identity to a type (T) sequence and/or >99.0% identity with a sequence that had not been typed, or (ii) >99.0% identity with >1 type (T) sequence in the database. We made presumptive species-level identifications based on a query sequence with >99.5% identity to a type sequence and only if that match was unambiguous (i.e. there was >99.5% identity with only 1 type sequence in the database).These species designations were made using reference sequences from the List of Prokaryotic names with Standing in Nomenclature (LSPN, ^48^, www. https://lpsn.dsmz.de/) (Table S1).

### Electroporation and GFP expression

Transformations of *P. zhaodongensis* (Pz15) were carried out using established protocols for electroporation of *Pseudomonas aeruginosa* ^49^ with no modifications. *P. zhaodongensis* (Pz15) isolates transformed with the GFP-expressing plasmid pMRP9 ^50^ were selected and propagated on LB agar containing 50 µg mL^-1^ carbenicillin.

### Genomic DNA purification for genome sequencing

Genomic DNA (gDNA) was isolated using a DNeasy® Blood and Tissue kit (Qiagen) with slight modifications to the manufacturer’s protocols. Briefly, bacteria were grown as required in 5 ml marine broth 2216 at 25 °C and 250 rpm overnight. Cells were then collected by centrifugation at 5000 X g for 10 min and washed twice with PBS. Pellets were suspended in 180 µl of ATL buffer (Qiagen) with 20 µl of proteinase K (Qiagen). Samples were incubated at 56 °C for 1 h, after which 20 µl RNase A (20 mg/ml) was added and left at room temperature for 5 min. Samples were then incubated for an additional 1 h at 56 °C and treated again with the addition of 20 µl RNase A for 5 min at room temperature before purification. DNA was then purified and eluted into 2 × 100 µl of elution buffer (EB, 10 mM Tris-HCl buffer, pH 7.5). Purity and quantity of gDNA were determined using absorbance measurements at 260 and 280 nm on a Nanodrop® and using the dsDNA HS assay kit (ThermoFisher) for the Qubit® 2.0. Sizes of purified DNA fragments were also assessed using standard protocols for analytical gel electrophoresis ^45^.

### Whole genome sequencing and assembly

Purified gDNA from *P. zhaodongensis* (Pz15), *Marinobacter adhaerens* HP15-B (Ma1), *Pseudoalteromonas shioyasakiensis* (Psh20), and *R. pomeroyi* DSS-3-B (Rp3) were sent to Genome Quebec for sequencing on the Pacific Biosciences RSII® or Sequel® platform. The gDNA libraries, which had an average insert size of 7 kb or larger, were prepared using the SMRTbell Express Template Prep Kit 2.0 (Pacific Biosciences). Genomes were sequenced with an estimated minimum 140-fold coverage. Contig assembly was carried out by the Canadian Center for Computational Genomics (C3G) using a Hierarchical Genome Assembly Process (HGAP) workflow ^51^. Sequencing data have been deposited with links to BioProject accession number PRJNA731154 in the National Center for Biotechnology Information (NCBI) BioProject database (https://www.ncbi.nlm.nih.gov/bioproject/).

### Genome annotation, comparative bioinformatics and genome visualizations

Genome annotation and comparative genomic analysis were conducted with Rapid Annotation using Subsystems Technology (RAST) ^42,52,53^. Annotations were made using the RASTtk annotation scheme with the Kmer dataset release 70 to annotate proteins and options to build a metabolic model. Genes predicted to encode putative proteins involved in fatty acid metabolism were identified by searching text annotations in the SEED platform ^54^ for International Union of Biochemistry and Molecular Biology (IUBMB) enzyme classification (EC) numbers retrieved from the Kyoto Encyclopedia of Genes and Genomes (KEGG) during annotation. Putative functions were corroborated by searching the NCBI Conserved Domain Database using the primary amino acid sequence of open reading frames identified in RAST. The complete *P. zhaodongensis* (Pz15), *M. adhaerens* HP15-B (Ma1), *P. shioyasakiensis* (Psh20), and *R. pomeroyi* DSS-3-B (Rp3) genomes were deposited in NCBI Genbank with accessions CP076683.1/CP076684.1, CP076686.1, CP076681.1/CP076682.1, and CP076685.1, respectively, corresponding to BioSamples SAMN19272545, SAMN19272548, SAMN19272546, and SAMN19272547, respectively. Visualizations were generated using ChemDraw Professional (Perkin-Elmer), and Illustrator Creative Cloud 2018 (Adobe).

### Molecular phylogenetics

The 16s rRNA gene sequences (Table S1) were aligned using MUSCLE ^55^. An approximately-maximum-likelihood phylogenetic tree was inferred from the alignment using the FastTree 2 ^56^ plug-in with default settings and visualization tools in Geneious Prime 2020.

### Mathematical model of vertical flux

We developed a mathematical model to explore how microbial interactions, via their impact on the rate of degradation of particles containing lipids and the time for initiation of their degradation, control the shape of the attenuation of the vertical flux of lipid with depth in the ocean. We assumed that degradation of the lipid droplet phase in lipid-ballast particles proceeds via a pseudo-first-order rate law, where the absolute rate of lipid degradation is dependent on the typical exposed lipid droplet surface area. The model considers spherical particles containing the non-degradable ballast and degradable lipid, present at a fixed universal initial ratio of volumes, generated at a reference depth *z*_0_ = 100 m, taken as representative of the bottom of the euphotic zone. The size distribution of these particles matches environmental values. For each initial radius of particles from this distribution, we model lipid droplet degradation based on our observed experimental degradation rates and particle sinking dynamics following Stokes law to obtain the particle concentration and lipid content per particle as a function of depth. By integrating the contributions of each particle size-class, we then compute the total vertical flux of lipids as a function of depth, and thus quantify the attenuation of this flux due to microbial degradation. This model, based on individual particle dynamics, is similar to recent approaches to capture vertical flux attenuation ^2,3,36^, but differs from previous approaches because it considers two distinct phases within each particle, namely an lipid droplet phase and a ballast phase. A complete description of the model is in Supplementary. Computer codes are available upon request.

## Author contributions

L.B., U.A., B.A.S.V.M. and R.S. designed research. L.B., U.A.,Y.Y. and S.S. conducted microbiology and microscopy experiments. J.E.H, H.F.F. and B.A.S.V.M conducted chemical extractions and lipidomic experiments. H.A. and J.J.H. conducted molecular experiments. T.M. provided access to bacterial collections. L.B., U.A., J.E.H., D.P.L., F.J.P., V.I.F., H.F., H.A., J.J.H., S.S. and B.A.S.V.M analyzed and interpreted data. L.B., B.A.S.M. and R.S. provided funding acquisition, project administration, and resources. D.P.L, V.I.F., F.J.P. and B.A.S.V.M. developed modeling approaches. L.B., U.A., S.S., F.J.P., B.A.S.V.M. and R.S. wrote the paper.

## Acknowledgments

We thank R. Naisbit for help with editing the manuscript. We acknowledge the NCMA (Bigelow Laboratory, Maine, USA) for large-scale growth and harvest of *P. tricornutum* cells. We thank Tracy Mincer for the provision of bacterial isolates.

## Data availability

The datasets and representative videos generated and analysed during the study are available in the Supplementary Information of the paper.

## Code availability

The computer codes used during the study are available upon request.

