## Supplementary for "Microbial metabolic specificity controls pelagic lipid export efficiency"

### drives pelagic lipid remineralization

**This PDF file includes:**

Full description of the lipid vertical flux model

Figs. S1 to S9

Tables S1

Captions for Video S1 to S7

**Other Supplementary Materials for this manuscript include the following:**

Video S1 to S7

### Full description of the lipid vertical flux

We developed a theoretical model to explore the implications of the two major findings described here: 1) that microbial interactions affect the rate of lipid droplet degradation; and 2) that microbial interactions affect the time for initiation of degradation of lipid droplets.

#### *General principles*

The model considers lipid-containing spherical particles generated at a reference depth $z_0$ = 100 m, typically the bottom of the euphotic zone, at a known concentration. For each initial radius of these particles, we model lipid degradation and particle sinking as a function of biological parameters to obtain the particle concentration and lipid content per particle at any depth. By summing the contributions of each size-class, we can then compute the total lipid flux with depth, and thus quantify the attenuation of the lipid flux due to microbial degradation. We assume that degradation of lipid proceeds via a pseudo-first-order rate law, where the absolute rate of lipid degradation is dependent on the exposed surface area of the lipid phase in the particle. Such a model based on individual particle dynamics depending on initial size is similar to recent theoretical approaches to capture vertical flux attenuation (*1*, *2*), but differs from previous approaches because it considers two distinct phases within each particle, namely an lipid phase and a ballast phase.

#### *Particle degradation dynamics*

TAGs, the dominant component in lipid, are less dense than water, and yet particulate matter in the ocean nonetheless sinks (*3*). Thus, we paired ballast minerals with the lipid fraction of the particle to initiate sinking. With $r_{\mathrm{tot,0}}$ the initial radius at depth $z_0$ of a given lipid–ballast particle, the initial total volume of the particle is

$$V_{\mathrm{tot,0}} = \frac{4\pi}{3} r_{\mathrm{tot,0}}^3 \ . \qquad (1)$$

The initial volume fraction of ballast is defined as

$$\phi_0 = \frac{V_{\mathrm{bal,0}}}{V_{\mathrm{tot,0}}} \ , \qquad (2)$$

where $V_{\mathrm{bal,0}}$ is the initial volume of ballast in the particle. Since in our simplified model the particle is only made of lipid and ballast, the total initial volume $V_{\mathrm{tot,0}}$ is the sum of initial ballast and initial lipid volumes, that is $V_{\mathrm{tot,0}} = V_{\mathrm{bal,0}} + V_{\mathrm{lipid,0}}$ with $V_{\mathrm{lipid,0}}$ the initial lipid volume in the particle. Therefore, we can also write

$$V_{\text{lipid},0} = (1 - \phi_0)V_{\text{tot},0}\,. \tag{3}$$

In this model, we use a fixed universal initial volume fraction of ballast $\phi_0 = 0.125$, which corresponds to the ballast load resulting in a sinking speed of 50 m day$^{-1}$ for a lipid–ballast particle of diameter 200 μm in our modeling framework. The ballast volume, being non-degradable, is held constant through time as the particle descends, while the lipid volume $V_{\text{lipid}}(t)$ decreases with time due to microbial degradation. The total volume of the particle at time *t* of its descent is thus

$$V_{\text{tot}}(t) = V_{\text{bal},0} + V_{\text{lipid}}(t)\,. \tag{4}$$

In order to solve the model dynamically for the degradation of the lipid phase, an appropriate degradation law is adopted. As we observe in our experiments, bacteria colonize the outside of lipid droplets and do not penetrate into the droplets' interior, therefore degrading lipid from its exposed surface only. To implement this surface degradation, we consider a mass degradation rate $D(t)$ in g s$^{-1}$ that is proportional to the typical surface area of lipid phase exposed in a mixed particle, which we take as the surface area $A_{\text{lipid}}(t)$ of a sphere of volume $V_{\text{lipid}}(t)$, that is

$$A_{\text{lipid}}(t) = 4\pi\, r_{\text{lipid}}(t)^2, \tag{5}$$

with $r_{\text{lipid}}(t)$ the equivalent spherical radius of the lipid phase given by

$$r_{\text{lipid}}(t) = \left[\frac{3\, V_{\text{lipid}}(t)}{4\,\pi}\right]^{\frac{1}{3}}. \tag{6}$$

Moreover, we incorporate our experimental observation that degradation starts after a delay in time $t_{\text{delay}}$ dependent on microbial interactions on the particle (Fig. 3, main text). This results in a mass degradation rate

$$D(t) = \begin{cases} 0 & \text{for} \quad t < t_{\text{delay}} \\ -\,k_A A_{\text{lipid}}(t) & \text{for} \quad t > t_{\text{delay}} \end{cases}, \tag{7}$$

with $k_A$ a mass degradation rate per unit surface area, a parameter of the model. Note that when the lipid phase is entirely degraded, $A_{\text{lipid}} = 0$ and therefore degradation in the model stops.

Relevant values for the lipid droplet degradation rate constant $k_A$ were determined from the degradation rates we observed experimentally (Fig. 3, main text) as

$$k_A = \frac{D_{\text{observed}}}{A_{\text{lipid}} \times f_{\text{coverage}}}, \quad (8)$$

where $D_{\text{observed}}$ is the observed degradation rate in g $s^{-1}$, $A_{\text{lipid}}$ is the observed total areas of the droplets spotted on the glass vials used in the experiments in $m^2$, and $f_{\text{coverage}}$ accounts for the patchiness of those droplet spots by giving the fraction of the drop areas effectively covered in lipid. The estimates of $D_{\text{observed}}$ ($n$ = 39), $A_{\text{lipid}}$ ($n$ = 20), and $f_{\text{coverage}}$ ($n$ = 40) were not collected in a manner that allowed us to calculate $k_A$ for discrete droplets. Therefore, a Monte-Carlo resampling simulation was conducted using the observed distribution of these three values to calculate a distribution of degradation rate constants (Suppl. Fig. S8). This distribution shows that $k_A$ values are typically in the range $2 \times 10^{-6}$ g $m^{-2}$ $s^{-1}$ to $1 \times 10^{-4}$ g $m^{-2}$ $s^{-1}$, which is the range adopted here.

Finally, the mass degradation rate $D(t)$ described above is coupled to the rate of change of the volume of lipid in the particle $V_{\text{lipid}}(t)$ via the density of the lipid phase $\rho_{\text{lipid}}$, that is

$$\frac{dV_{\text{lipid}}}{dt} = \frac{D(t)}{\rho_{\text{lipid}}}. \quad (9)$$

The density of the lipid phase $\rho_{\text{lipid}}$ was chosen to match the density of one of the prevalent TAGs in the phytoplankton lipid extract used in our experiments, tripalmitin ($8.8 \times 10^5$ g $m^{-3}$). Note that TAG density only increases by about $0.1 \times 10^5$ g $m^{-3}$ (i.e., ≈ 1%) when compressed to 10 MPa (≈ 1,000 dbar), (*4*) and therefore, the model ignores the effects of pressure-induced density changes on lipid droplets during their descent.

#### *Particle sinking dynamics*

Particles in the model sink through the water column while they are being degraded. Their time-varying vertical position $z(t)$ is obtained by integrating their sinking speed $S(t)$, which we assume is a function of their total size and density according to Stokes sedimentation law, that is

$$S(t) = \frac{dz}{dt} = \frac{2\,g}{9\,\mu}\,[\rho_{\text{tot}}(t) - \rho_{\text{sw}}]\; r_{\text{tot}}^2(t), \quad (10)$$

where $g$ is gravitational acceleration (9.8 m $s^{-2}$), $\mu$ is the dynamic viscosity of seawater (1.40 g $m^{-1}$ $s^{-1}$ for seawater at 10 ºC), $\rho_{\text{sw}}$ is the density of seawater (which is held constant for simplicity; $1.027 \times 10^6$ g m$^{-3}$), and $r_{\mathrm{tot}}(t)$ is the radius of the entire particle, that is

$$r_{\mathrm{tot}}(t) = \left[\frac{3\, V_{\mathrm{tot}}(t)}{4\,\pi}\right]^{\frac{1}{3}}. \qquad (11)$$

In Equation (10), the average density of the particle $\rho_{\mathrm{tot}}(t)$ takes into account both the lipid phase and the ballast phase, and is defined as

$$\rho_{\mathrm{tot}}(t) = \frac{V_{\mathrm{lipid}}(t)}{V_{\mathrm{lipid}}(t)+V_{\mathrm{bal,0}}}\ \rho_{\mathrm{lipid}} + \frac{V_{\mathrm{bal,0}}}{V_{\mathrm{lipid}}(t)+V_{\mathrm{bal,0}}}\ \rho_{\mathrm{bal}}\ . \qquad (12)$$

In this expression, $\rho_{\mathrm{bal}}$ is the density of the ballast phase, chosen here to match the density of a 50/50 mix of opal/calcium carbonate ($2.35 \times 10^6$ g m$^{-3}$), and $\rho_{\mathrm{lipid}}$ is the density of the lipid phase as described above.

***Quantification of transfer efficiency for single size class and over particle distribution***

By solving the coupled equations of lipid degradation dynamics (Equation (9)) and sinking dynamics (Equation (10)), one can obtain for each chosen particle of initial radius $r_{\mathrm{tot,0}}$ the time evolution of its associated lipid volume $V_{\mathrm{lipid}}(t)$ and its position $z(t)$ (see section "Analytical solution" below for details).

We assume the initial range of radii is defined by $r_l < r_{\mathrm{tot,0}} < r_g$, with $r_l$ the minimum particle radius at $z_0$ and $r_g$ the maximum particle radius at $z_0$. We assume a distribution of particle initial radii $N(r_{\mathrm{tot,0}})$ over that range, so that the number of particles of radii between $r_{\mathrm{tot,0}}$ and $r_{\mathrm{tot,0}} + \mathrm{d}r_{\mathrm{tot,0}}$ per unit volume seawater at depth $z_0$, is given by $N(r_{\mathrm{tot,0}})\, \mathrm{d}r_{\mathrm{tot,0}}$. Specifically, we consider particles with initial radii from $r_l$ = 25 µm to $r_g$ = 250 µm (*5*), following a decreasing power law distribution so that

$$N\left(r_{\mathrm{tot,0}}\right) = \alpha\, r_{\mathrm{tot,0}}^{\beta}, \qquad (13)$$

with β = - 2.965 as observed (*5*). Such a distribution means that particles of radius 25 µm are initially three orders of magnitude more abundant than particles of radius 250 µm. In equation (13), $\alpha$ is a constant characteristic of the total concentration of particles. As we will only consider relative values of lipid fluxes, its exact value does not influence the results of our analysis of the model.

As particles shrink in our model, their sinking speed decreases. We assume a steady state of particle concentration over the water column, and therefore a constant number vertical flux (in particles per unit area per unit time) with depth. Under this hypothesis of steady state, the decrease in speed is balanced by an increase in local concentration per unit volume seawater. Specifically, the local concentration $C$ at depth $z$ of particles with initial radius (at depth $z_0$) comprised between $r_{\mathrm{tot,0}}$ and $r_{\mathrm{tot,0}}$ + $\mathrm{d}r_{\mathrm{tot,0}}$ is

$$C(z, r_{\mathrm{tot,0}}) = N(\mathrm{r}_{\mathrm{tot,0}})\, \mathrm{d}r_{\mathrm{tot,0}} \frac{S_0(\, r_{\mathrm{tot,0}})}{S[t_{\mathrm{s}}(z, r_{\mathrm{tot,0}})]}, \tag{14}$$

where $S_0(r_{\mathrm{tot,0}})$ is the initial speed of these particles at depth $z_0$, and $t_{\mathrm{s}}(z,\, r_{\mathrm{tot,0}})$ is the time necessary for a particle of initial radius $r_{\mathrm{tot,0}}$ to sink to depth $z$. Therefore, $S\,[t_{\mathrm{s}}(z,\, r_{\mathrm{tot,0}})]$ is the sinking speed of particles of initial radius $r_{\mathrm{tot,0}}$ when they reach depth $z$, and the concentration of particles increases in a manner inversely proportional to the decreasing sinking speed.

We can then at any depth $z$ calculate the lipid partial transfer efficiency $T_{\mathrm{p}}(z,\, r_{\mathrm{tot,0}})$ for particles of a given initial radius $r_{\mathrm{tot,0}}$, as the ratio of the vertical lipid flux at depth $z$ over the vertical lipid flux associated with these particles at initial depth $z_0$. With these fluxes being the product of lipid volume per particle, particle concentration and vertical speed, we have

$$T_{\mathrm{p}}(z, r_{\mathrm{tot,0}}) = \frac{V_{\mathrm{lipid}}[t_{\mathrm{s}}(z, r_{\mathrm{tot,0}})]\;\; C(z, r_{\mathrm{tot,0}})\;\; S[t_{\mathrm{s}}(z, r_{\mathrm{tot,0}})]}{V_{\mathrm{lipid,0}}(r_{\mathrm{tot,0}})\;\; C(z_0, r_{\mathrm{tot,0}})\;\; S_0(\, r_{\mathrm{tot,0}})}\ , \tag{15}$$

which using the expression for the concentration in equation (14) simplifies to

$$T_{\mathrm{p}}(z, r_{\mathrm{tot,0}}) = \frac{V_{\mathrm{lipid}}[t_{\mathrm{s}}(z, r_{\mathrm{tot,0}})]}{V_{\mathrm{lipid,0}}(r_{\mathrm{tot,0}})}\ . \tag{16}$$

This is simply the ratio of the single-particle lipid volume when reaching depth $z$ at time $t_{\mathrm{s}}(z,\, r_{\mathrm{tot,0}})$ over the single-particle initial lipid volume $V_{\mathrm{lipid,0}}(r_{\mathrm{tot,0}})$ for this choice of initial radius.

The attenuation of flux for each initial radius class will sum up to give the attenuation with depth of the total lipid flux associated with all particles considered. To characterize this total attenuation, we can first compute the total vertical flux of lipid $F_{\mathrm{lipid}}$ at any depth $z$, which is obtained by integration over the range of the initial radius as

$$F_{\text{lipid}}(z) = \int_{r_\text{l}}^{r_\text{g}} V_{\text{lipid}}\left[t_\text{s}\left(z, r_{\text{tot},0}\right)\right]\ C\left(z, r_{\text{tot},0}\right)\ S\left[t_\text{s}\left(z, r_{\text{tot},0}\right)\right]\ \mathrm{d}r_{\text{tot},0}\ , \tag{17}$$

which using equation (14) simplifies to

$$F_{\text{lipid}}(z) = \int_{r_\text{l}}^{r_\text{g}} V_{\text{lipid}}\left[t_\text{s}\left(z, r_{\text{tot},0}\right)\right]\ N\left(r_{\text{tot},0}\right)\ S_0\left(r_{\text{tot},0}\right)\ \mathrm{d}r_{\text{tot},0}\ . \tag{18}$$

The attenuation of the vertical flux of lipid over the entire range of initial particle radii can be characterized at a given depth $z$ by the transfer efficiency $T(z)$ with respect to initial depth $z_0$ (*6*, *7*), which is the ratio of flux at depth $z$ over initial flux, that is

$$T(z) = \frac{F_{\text{lipid}}(z)}{F_{\text{lipid}}(z_0)}\ . \tag{19}$$

We run the model using bacterial degradation rate constants $k_A$ spanning the range found in our experiments ($2 \times 10^{-6}$ g m$^{-2}$ s$^{-1}$ to $1 \times 10^{-4}$ g m$^{-2}$ s$^{-1}$), while also examining the effects of different delays in the start of degradation $t_{\text{delay}}$, chosen to span our experimental observations (0, 6, 12, 24 hours). The dynamics of degradation and sinking for a given initial particle radius are directly obtained from the analytical solution described in the section below, and the partial transfer efficiencies $T_\text{p}(z, r_{\text{tot},0})$ and total transfer efficiency $T(z)$ are evaluated numerically using a custom Python code. We consider in particular the transfer efficiencies at depth $z_0$ + 100 m and depth $z_0$ + 900 m (Fig. S8).

#### *Analytical solution of single particle sinking and degradation dynamics*

In this section, we derive an analytical solution of the coupled problem of sinking and degradation described above for any given initial radius $r_{\text{tot},0}$.

We can rewrite the lipid degradation dynamics of equation (9) as

$$4\,\pi\, r_{\text{lipid}}(t)^2 \frac{dr_{\text{lipid}}}{dt} = \begin{cases} 0 & \text{for } t < t_{\text{delay}} \\ -\dfrac{k_A}{\rho_{\text{lipid}}} 4\,\pi\, r_{\text{lipid}}(t)^2 & \text{for } t > t_{\text{delay}} \end{cases} \tag{20}$$

using the degradation rate expression in equation (7). Introducing the Heaviside function $H(t)$ equal to 0 for $t < 0$ and 1 for $t > 1$, we can write the solution of equation (20) as

$$r_{\text{lipid}}(t) = \begin{cases} r_{\text{lipid,0}} - \dfrac{k_A}{\rho_{\text{lipid}}} t \times H(t - t_{\text{delay}}) & \text{for } \; t < t_{\text{delay}} + t_{\text{lipid}} \\ 0 & \text{for } \; t > t_{\text{delay}} + t_{\text{lipid}} \end{cases} \tag{21}$$

with $t_{\text{lipid}} = r_{\text{lipid,0}}\ \rho_{\text{lipid}} / k_A$ the time necessary to degrade the entire lipid phase in the case with no delay ($t_{\text{delay}}$ = 0 h). From this expression, one can directly obtain the analytical solution for the time evolution of the lipid volume as $V_{\text{lipid}}(t) = 4\pi\ r_{\text{lipid}}(t)^3/3$. One can also write analytically the total radius of the particle as

$$r_{\text{tot}}(t) = \left[r_{\text{lipid}}(t)^3 + r_{\text{bal,0}}^3\right]^{\frac{1}{3}} \tag{22}$$

and use equation (21) to replace $r_{\text{lipid}}(t)$ by its analytical expression to get an analytical expression for $r_{\text{tot,t}}$.

We can then obtain an analytical solution of the sinking speed by noticing that combining equations (10) for the definition of sinking speed, (12) and (22) results in

$$S(t) = \frac{2\,g}{9\,\mu}\left[r_{\text{lipid}}(t)^3(\rho_{\text{lipid}} - \rho_{\text{sw}}) + r_{\text{bal,0}}^3(\rho_{\text{bal}} - \rho_{\text{sw}})\right]\left[r_{\text{lipid}}(t)^3 + r_{\text{bal,0}}^3\right]^{-\frac{1}{3}} \tag{23}$$

and using equation (21) to replace $r_{\text{lipid}}(t)$ by its analytical expression.

We integrate equation (23) with respect to time to obtain the depth with time $z(t)$. We introduce the notations

$$\theta(t) = \frac{r_{\text{lipid}}(t)}{r_{\text{tot}}(t)} \quad \text{with} \quad \theta(0) = \theta_0 = \frac{r_{\text{lipid,0}}}{r_{\text{tot,0}}} \tag{24}$$

for the ratio of equivalent lipid radius to total radius, for which an analytical expression can then be obtained by combining equations (21) and (22). We also introduce the notations $\Delta\rho_{\text{lipid}} = \rho_{\text{lipid}} - \rho_{\text{sw}}$ and $\Delta\rho_{\text{bal}} = \rho_{\text{bal}} - \rho_{\text{sw}}$. We find that in the no-delay case ($t_{\text{delay}} = 0$), the sinking speed given by equation (23) can be integrated explicitly to give an analytical expression for the position with time

$$z_{\text{no delay}}(t) = z_0 + \frac{\rho_{\text{lipid}} g}{81\, k_A \mu}\Bigg\{ \; 6\,\Delta\rho_{\text{lipid}}\big(\theta_0\, r_{\text{tot},0}^3 - \theta(t) r_{\text{tot}}(t)^3\big)$$
$$+ (3\,\Delta\rho_{\text{bal}} - \Delta\rho_{\text{lipid}}) r_{\text{bal},0}^3 \left[ 2\sqrt{3}\arctan\left(\frac{1+2\theta_0}{\sqrt{3}}\right) - 2\sqrt{3}\arctan\left(\frac{1+2\theta(t)}{\sqrt{3}}\right) - 2\log\left(\frac{1-\theta_0}{1-\theta(t)}\right) + \log\left(\frac{1+\theta_0+\theta_0^2}{1+\theta(t)+\theta(t)^2}\right)\right]\Bigg\}. \tag{25}$$

This solution is valid for $t < t_{\text{lipid}}$, after which time all the lipid is consumed and the particle consists only of non-degradable ballast and continues sinking at constant speed.

Using this reference solution for the no-delay case, one can write a general expression for the depth $z(t)$ reached by a given particle with a delay $t_{\text{delay}}$ as

$$z(t) = \begin{cases} S_0 t & \text{for } t < t_{\text{delay}}\,, \\ S_0 t_{\text{delay}} + z_{\text{no delay}}(t - t_{\text{delay}}) & \text{for } t_{\text{delay}} < t < t_{\text{delay}} + t_{\text{lipid}}, \\ S_0 t_{\text{delay}} + z_{\text{no delay}}(t_{\text{lipid}}) + S_{\text{bal}}(t - t_{\text{delay}} - t_{\text{lipid}}) & \text{for } t > t_{\text{delay}} + t_{\text{lipid}} \end{cases} \tag{26}$$

with

$$S_0 = \frac{2\,g}{9\,\mu}\,(\rho_{\text{tot},0} - \rho_{\text{sw}})\; r_{\text{tot},0}^2 \tag{27}$$

the initial constant sinking speed before the onset of degradation and

$$S_{\text{bal}} = \frac{2\,g}{9\,\mu}\,(\rho_{\text{bal}} - \rho_{\text{sw}})\; r_{\text{bal},0}^2 \tag{28}$$

the final constant sinking speed once the lipid phase is degraded.

Finally, we can numerically invert equation (26) to obtain the time $t_s(z, r_{\text{tot},0})$ at which a particle of initial radius $r_{\text{tot},0}$ will reach depth $z$, and compute the associated lipid volume and sinking speed using equations (21) and (23) to then calculate the transfer efficiencies, as presented above.

#### *Estimating bacterial lipid degradation*

In order to estimate the quantity of lipid degradation attributable to bacterial activity, changes in lipid droplet content were first calibrated against the changes in lipid autofluorescence. This was achieved sequentially fitting a known empirical relationship between lipid autofluorescence quenching and chlorophyll (Fig. S4A) and then a volume-based relationship between chlorophyll and lipid content in a droplet (Fig. S4B). The combination results in a physically-grounded approximately linear relationship between lipid autofluorescence quenching and lipid content in a droplet (Fig. S4C).

Following Kelly and Porter (*8*), the fluorescent yield from lipid autofluorescence follows the following formula:

$$\varphi_n = \frac{1}{1 + \left( {C}/{C_{0.5}} \right)^2} \qquad (29)$$

where $\varphi_n$ is the normalized fluorescence, $C$ is the chlorophyll concentration, and $C_{0.5}$ is a constant corresponding to the chlorophyll concentration resulting in half of the maximum fluorescence. In practice, the initial state of the lipid droplets does not correspond to such a low concentration of chlorophyll that $C/C_{0.5}$ is approximately zero. Therefore, we fit the experimental data for both $C_{0.5}$ (14.72 mM) and a constant of proportionality (1.885) that renormalizes the fluorescence intensity.

The volume of a lipid droplet is composed of lipids, chlorophyll, and other miscellaneous material. Of all the contents, we assume that only the lipids are degraded by bacteria and that they initially represent a large fraction of the droplet. As a result, the chlorophyll concentration (inversely proportional to droplet volume) and the total lipid content were modeled to have a simple inverse relationship:

$$L = \frac{a}{C}$$

Note that the lipid content $L$ is a total quantity over the droplet (nmol C), whereas the chlorophyll $C$ is a concentration (mol/L). The constant was found to be 1.1473. This simplified relationship does not hold when the total volume of lipid becomes comparable to or smaller than the volume of non-lipid content of the droplet. This corresponds to the right extreme of Fig. S4B. However, we felt that there was not enough data in this regime to parameterize a more complex model, and the lipid degradation metrics of interest would occur during situations where the lipid content dominated the droplets.

### Supplementary Figures

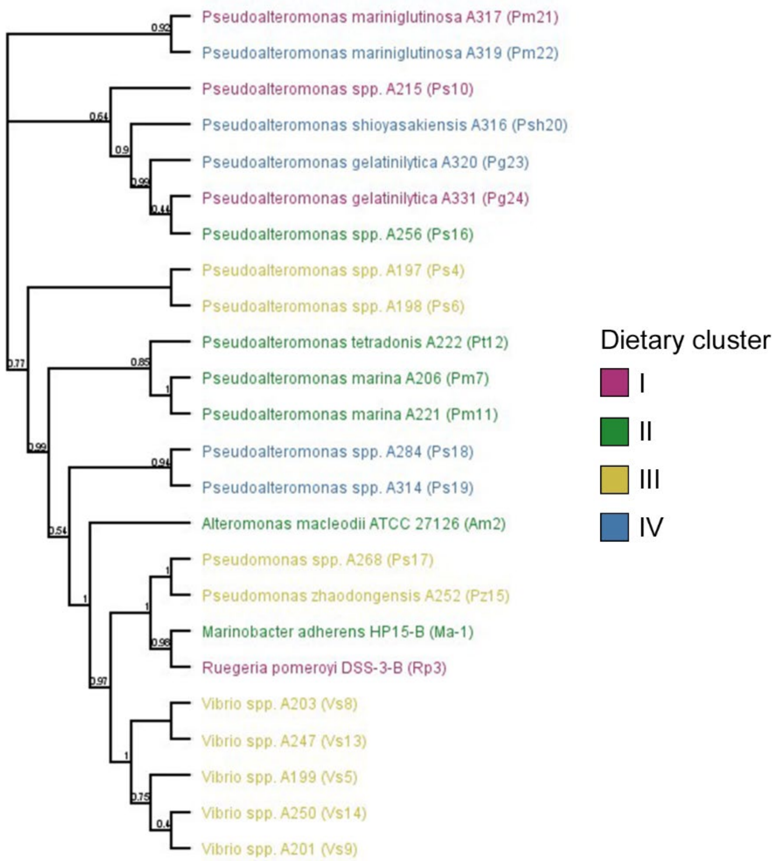


**Fig. S1. Phylogenetics cannot be used to predict the lipid degradation preferences of marine bacterial species.** Maximum-likelihood molecular phylogeny cladogram of 16S rRNA gene sequences for the 24 marine bacterial species investigated in this study. Genus and species are color coded according to their lipid dietary cluster. The phylogeny was constructed using FastTree 2 and values represent local support values based on the Shimodaira–Hasegawa (SH) test (*9*).

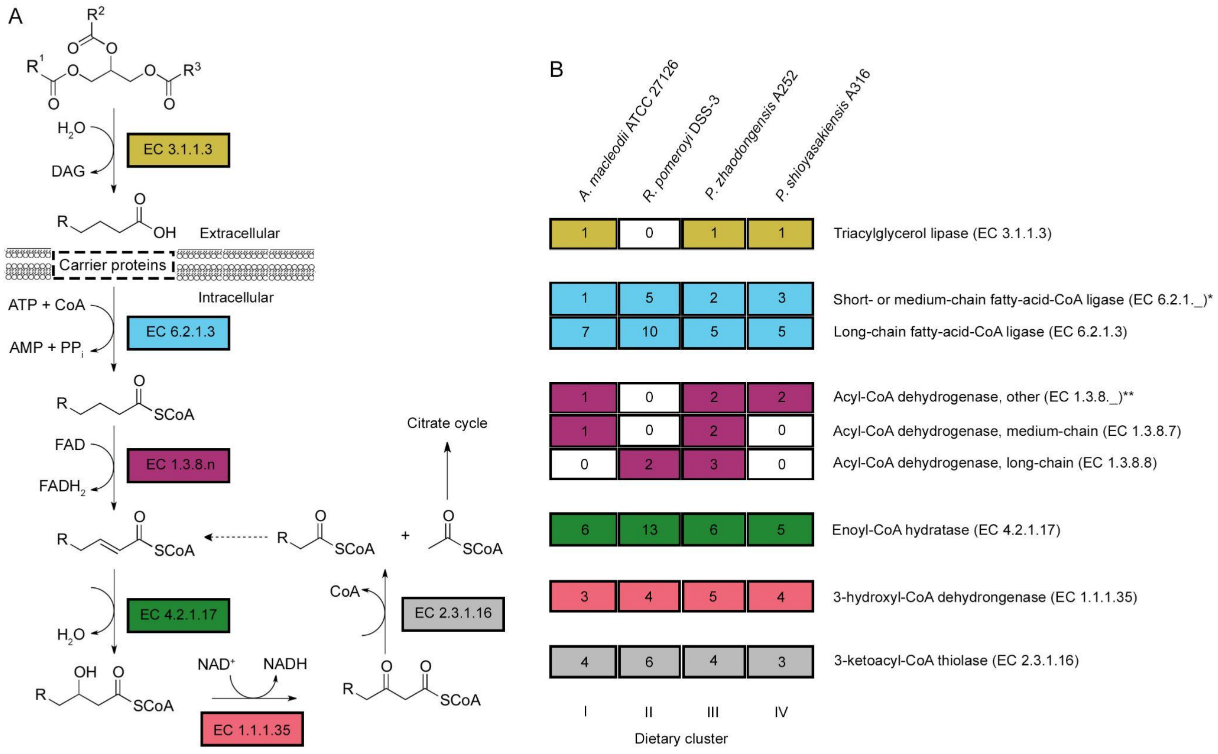


| Enzyme | I | II | III | IV |
|---|---|---|---|---|
| Triacylglycerol lipase (EC 3.1.1.3) | 1 | 0 | 1 | 1 |
| Short- or medium-chain fatty-acid-CoA ligase (EC 6.2.1._)* | 1 | 5 | 2 | 3 |
| Long-chain fatty-acid-CoA ligase (EC 6.2.1.3) | 7 | 10 | 5 | 5 |
| Acyl-CoA dehydrogenase, other (EC 1.3.8._)** | 1 | 0 | 2 | 2 |
| Acyl-CoA dehydrogenase, medium-chain (EC 1.3.8.7) | 1 | 0 | 2 | 0 |
| Acyl-CoA dehydrogenase, long-chain (EC 1.3.8.8) | 0 | 2 | 3 | 0 |
| Enoyl-CoA hydratase (EC 4.2.1.17) | 6 | 13 | 6 | 5 |
| 3-hydroxyl-CoA dehydrongenase (EC 1.1.1.35) | 3 | 4 | 5 | 4 |
| 3-ketoacyl-CoA thiolase (EC 2.3.1.16) | 4 | 6 | 4 | 3 |

**Fig. S2. Marine bacteria from different dietary clusters possess distinct repertoires of putative enzymes for catabolic lipid metabolism.** (A) Predicted pathway for triacylglycerol metabolism and β-oxidation in bacteria (adapted from the KEGG database, EC00071). Enzymes catalyzing the indicated reaction are color coded by number in the International Union for Biochemistry and Molecular Biology (IUBMB) Enzyme Classification scheme. (B) Presence/absence analysis of putative genes involved in triacylglycerol metabolism and β-oxidation in the genomes of marine bacterial species representing each lipid dietary cluster. The values in the matrix indicate the number of paralogs of each gene encoding the predicted enzymatic function in the genome, color coded by IUBMB Enzyme Classification. An asterisk (*) denotes putative acyl-coA ligases that may have specificity for short-chain (EC 6.2.1.1, EC 6.2.1.17) or medium-chain (EC 6.2.1.2) acyl substrates. A double asterisk (**) denotes putative acyl-CoA dehydrogenases that may have specificity for short-chain acyl substrates (EC 1.3.8.1) or for which an accurate substrate-length prediction cannot be made. . Putative enzymes for long-chain substrates are predicted to have specificity for acyl groups with 13 to 21 carbon atoms. Paralogs in the same category have likely diverged to accommodate acyl chains of different lengths, leading to the prediction that numbers of putative enzymes in each category correspond to a breadth in digestive capacity for different fatty acyl substrates.

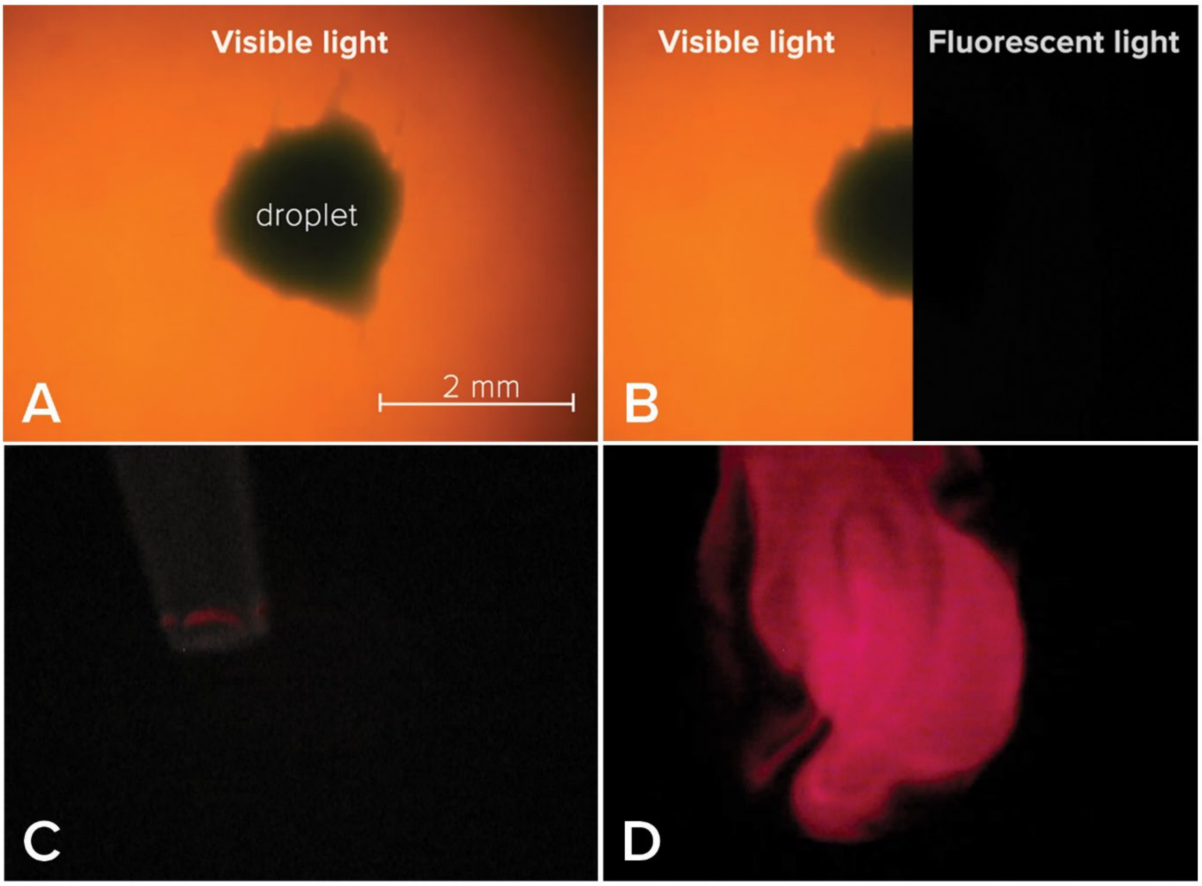


**Fig. S3. Video sequence of the concentration-dependent quenching of chlorophyll fluorescence within a lipid droplet**. (A) A droplet containing highly concentrated chlorophyll viewed under transmitted visible light. (B) Split screen view of the droplet showing that under epifluorescence (Ex.: 628/40 nm HBW, Em.: 692/40 nm HBW), the chlorophyll does not emit fluorescent light due to chlorophyll-chlorophyll self-quenching. (C) a glass pipette containing triacylglycerol (trioctanoylglycerol) was positioned above the same droplet; note that the background levels of this panel were altered slightly to make the pipette more visible, but remain unaltered in the accompanying video (Video S4). (D) Additional triacylglycerol was dispensed from the pipette into the droplet diluting the chlorophyll, which unquenched fluorescence.

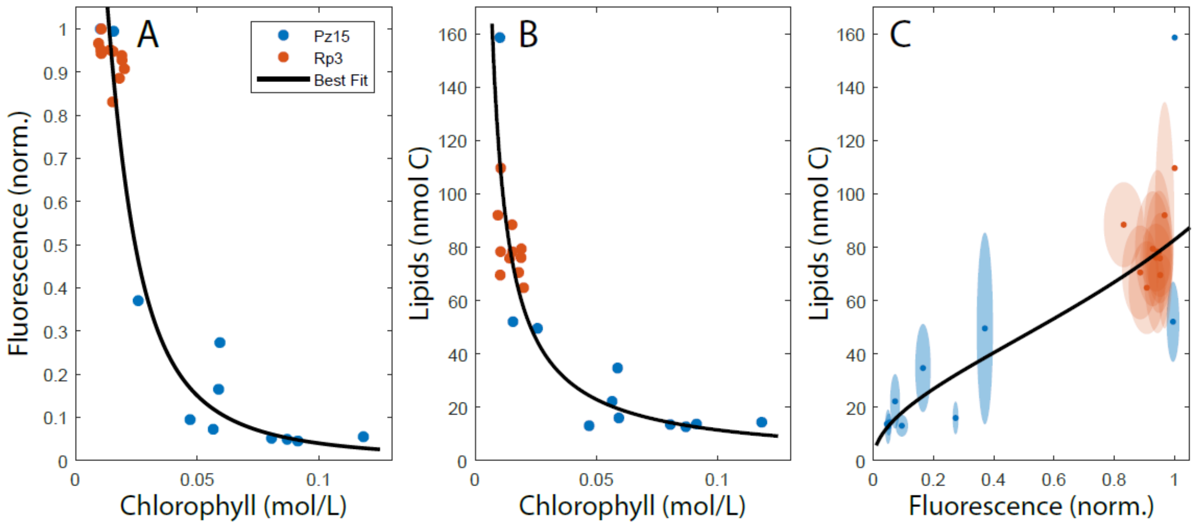


**Fig. S4. Calibration of the relation between autofluorescence and lipid content.** The parameters for a model of chlorophyll fluorescence quenching were identified by fitting a least-squares cost function to the average normalized fluorescence measurements (Fig. S5) and concurrent chlorophyll concentration measurements (A). Using data from the same droplets, parameters were obtained for a best-fit physical droplet model relating the average chlorophyll concentration and droplet lipid (B). The physical droplet model assumes that droplets are composed of a degradable lipid volume fraction, and a small fixed fraction of chlorophyll. As the lipids are degraded, the total droplet volume decreases proportionately, and the chlorophyll concentration correspondingly increases. Combined, these physically grounded relationships provide the calibration between droplet fluorescence and droplet lipid content (C). For the final composite calibration, the error ellipses with 1 standard deviation are shown, based on experiment replicates.

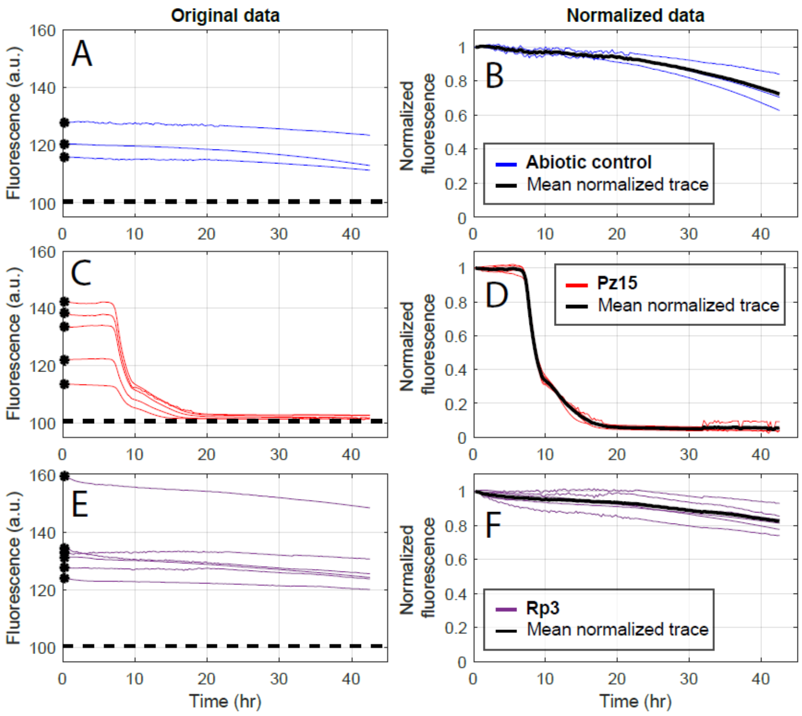


**Fig. S5. The normalization procedure used for lipid autofluorescence**. All fluorescence traces were normalized by first subtracting a fluorescence baseline (dashed line, A, C, E), and then dividing by the initial fluorescence (black points, A, C, E). Normalized fluorescence traces (B, D, F) all start at 1 as a result. The fluorescence baseline was determined to be the lower limit of the endpoint fluorescence for the strong degrader Pz15 (C). Subsequent co-culture experiments included a set of explicit lipid-free control wells that directly quantified the fluorescence baseline. The mean normalized fluorescence (black line, B, D, F) was used for calibration against lipid and chlorophyll measurements.

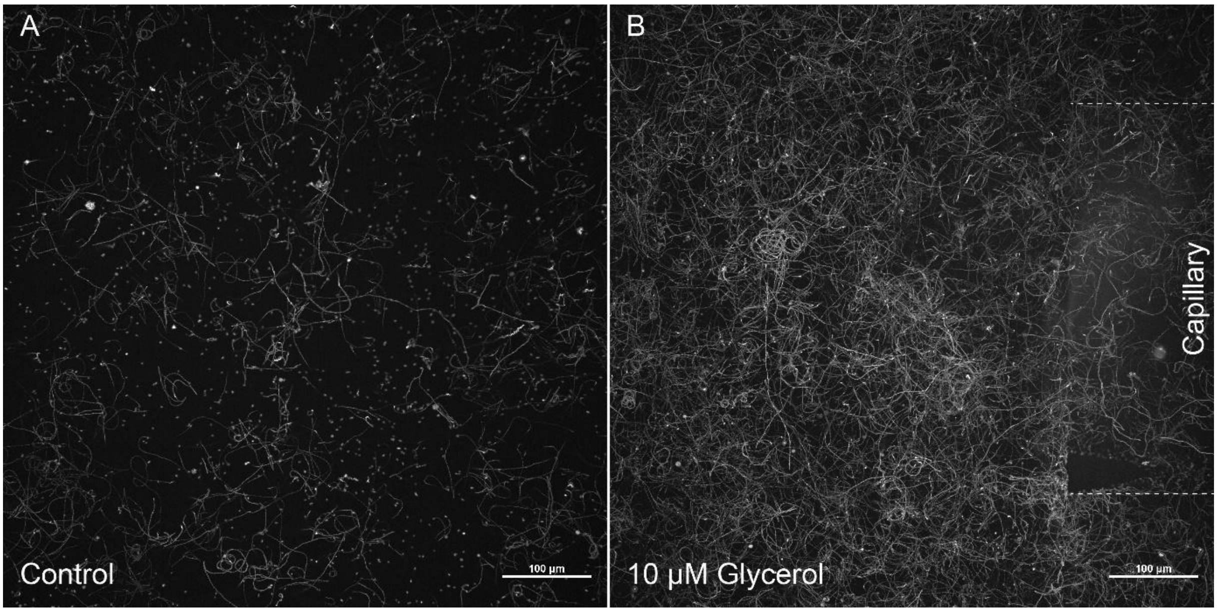


**Fig. S6. *Pseudomonas zhaodongensis* (Pz15) is positively chemotactic towards glycerol, the cleavage product released during TAG degradation**. (A) Swimming behavior of GFP-labeled Pz15 in f/2 medium. The image shows the maximum intensity projection of 500 frames recorded at 10 frames per second (fps). (B) Swimming behavior of Pz15 in the presence of a capillary filled with f/2 containing 10 µM glycerol. The location of the capillary is highlighted by dashed lines. The image shows 500 frames recorded at 10 fps after 15 min of incubation.

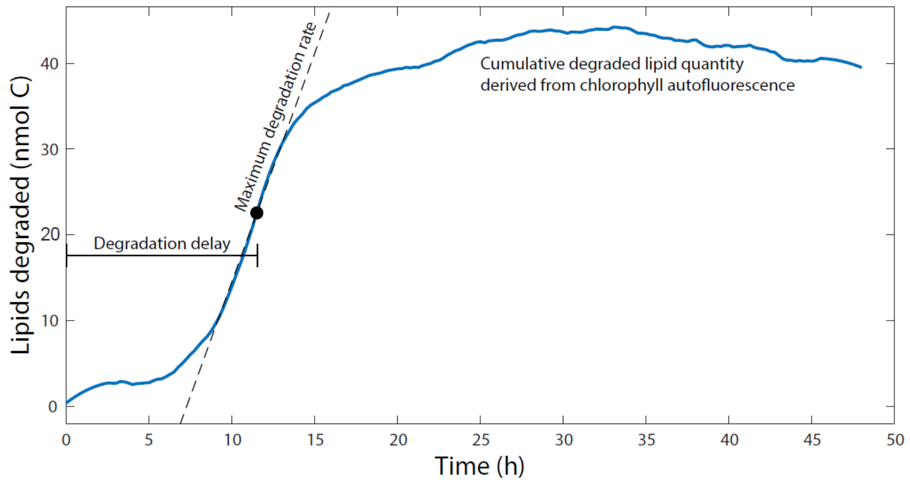


**Fig. S7. Example lipid degradation profile showing the parameters used to characterize breakdown**. The blue curve depicts the time course of lipid breakdown by bacteria, as derived from the lipid autofluorescence (Fig. S5) by comparing the reduction in lipid content in the presence of bacteria to that of droplet without bacteria present. The maximum degradation rate and degradation delay are shown, where the latter is the time until the point of maximum degradation as determined by smoothing with a Savitzky-Golay filter (5 hours, quadratic).

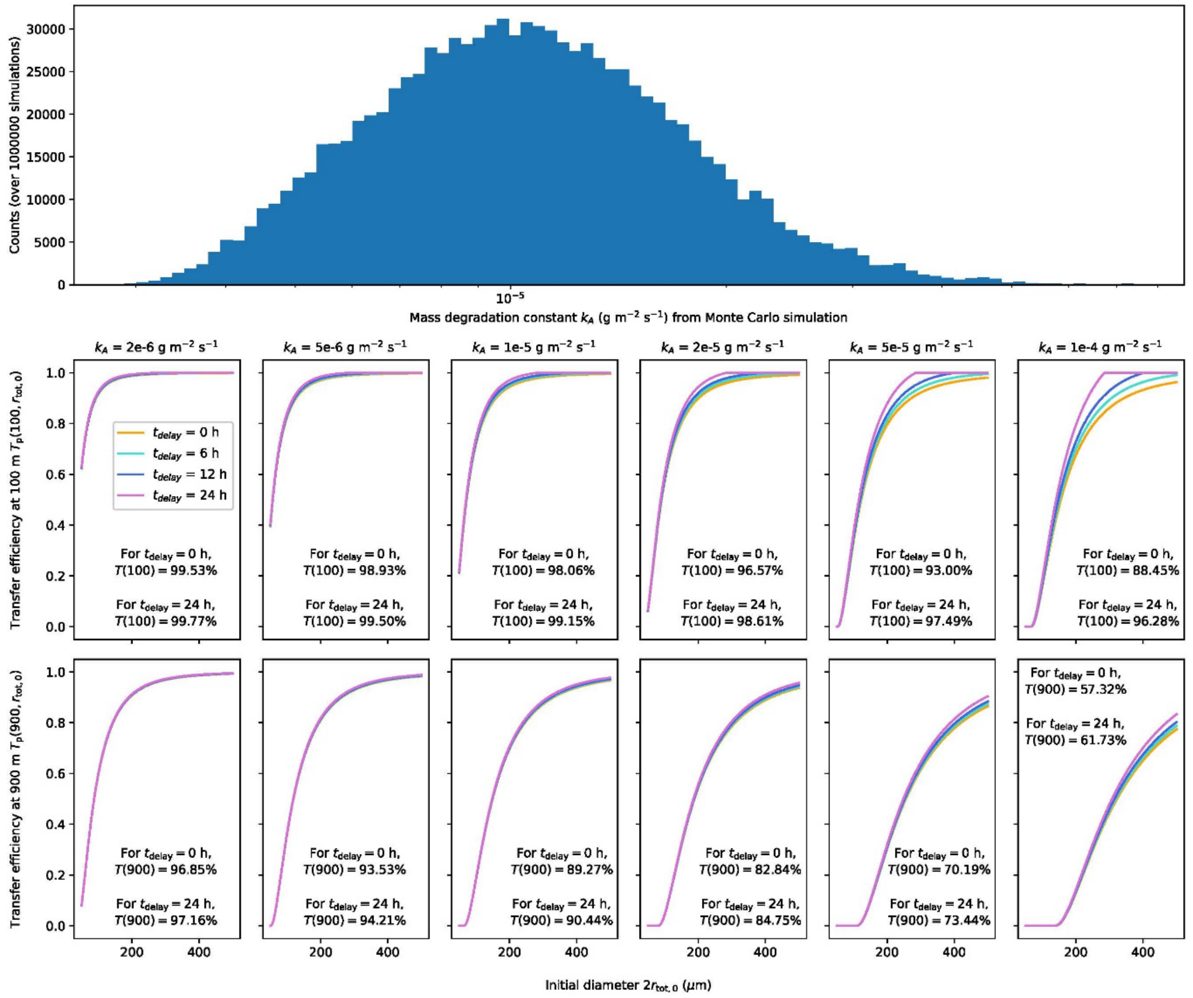


**Figure S8. Lipid transfer efficiency in the ocean is dependent on microbial interactions through effects on lipid degradation rate and degradation delay.** (**A**) Results of Monte-Carlo resampling simulation showing the distribution of possible degradation rate constants $k_A$ from our laboratory experiments (Fig. 3), based on random combinations of total degradations rates $D$, lipid droplet surface area $A_{lipid}$ and patchiness factor $f_{coverage}$ according to equation (8). (**B,C**) Plots of the transfer efficiency at $z$ = $z_0$ + 100 m (**B**) and $z$ = $z_0$ + 900 m (**C**). The $x$-axis shows the initial particle diameter ($2r_{tot,0}$) and the $y$-axis shows the partial transfer efficiencies $T_p(z, r_{tot,0})$. Results are shown for six values of the degradation rate constant ($k_A$) spanning those we observed in experiments. For each degradation rate, the total transfer efficiency $T(z)$ integrated over the entire range of initial particle radii between 25 μm and 250 μm is given for the no-delay case and for $t_{delay}$ = 24 h. The effects of different degradation delays on transfer efficiency is shown by different colors, with magenta representing 0 h delay.

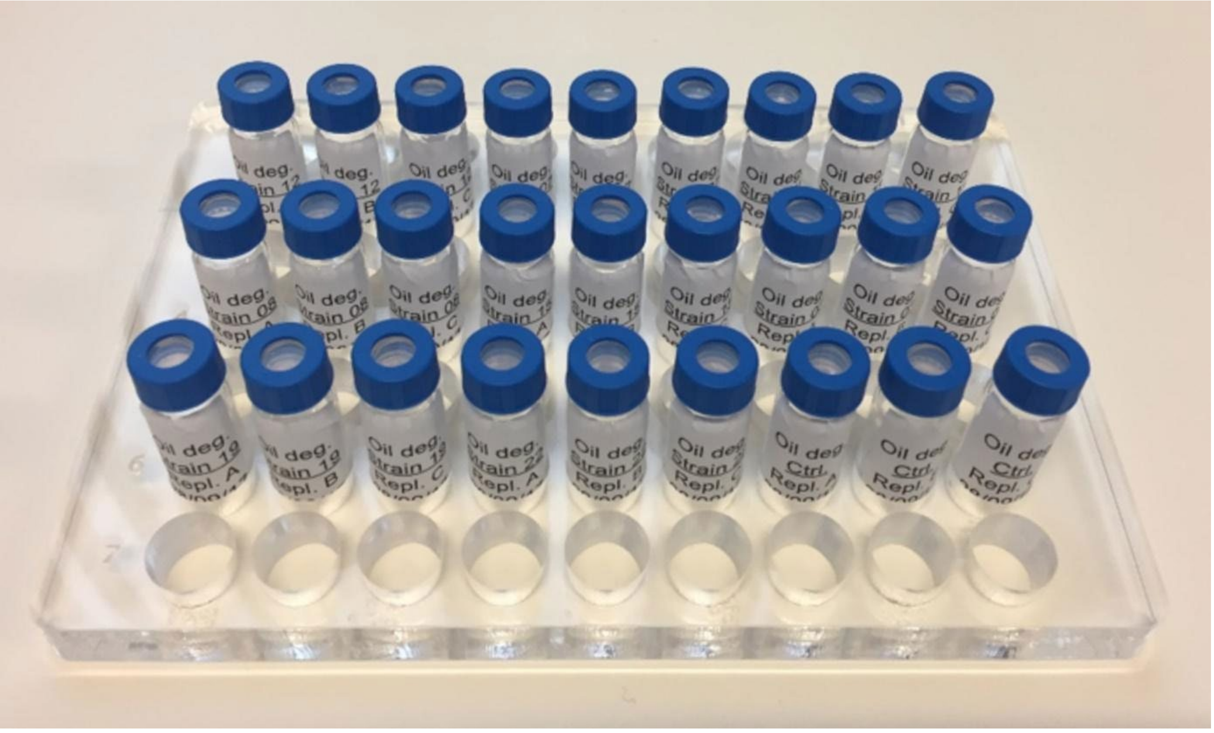


**Fig. S9. A microscopy-compatible acrylic holder for the simultaneous imaging of lipid droplets and lipidomic sampling**. This system enabled for the microscopic imaging of lipid droplets and bacteria through the bottom of LC-MS compatible vials. Following imaging, individual vials were removed at different time-intervals and chemically fixed for subsequent lipidomic analysis. As the removal of vials was done manually we noticed shifts in XY positions within fluorescent images. This required the development of an *in silico* jump correction to normalize signal shifts (Materials and Methods, Fig. S5)

**Supplementary Table 1.** Bacterial strains and plasmids used in this study. Names of isolates are followed by their abbreviated designators used in this study (Fig. 2).

| Strain or plasmid | Description[1] | Source |
|---|---|---|
| **Bacteria** | | |
| *Escherichia coli* NEB5α | Cloning strain | New England Biolabs |
| *Alteromonas macleodii* ATCC27126 (Am2) | Wild type | Oscar Sosa, E. F. DeLong lab |
| *Marinobacter adhaerens* HP15-B (Ma1) | Wild type | M. Ullrich, Jacobs University |
| *Pseudoalteromonas gelatinilytica* (Pg23) | Wild type; marine isolate | This study |
| *Pseudoalteromonas sp.* (Ps10) | Wild type; marine isolate | This study |
| *Pseudoalteromonas marina* (Pm7) | Wild type; marine isolate | This study |
| *Pseudoalteromonas marina* (Pm11) | Wild type; marine isolate | This study |
| *Pseudoalteromonas mariniglutinosa* (Pm21) | Wild type; marine isolate | This study |
| *Pseudoalteromonas mariniglutinosa* (Pm22) | Wild type; marine isolate | This study |
| *Pseudoalteromonas shioyasakiensis* (Psh20) | Wild type; marine isolate | This study |
| *Pseudoalteromonas tetradonis* (Pt12) | Wild type; marine isolate | This study |
| *Pseudoalteromonas* spp. (Ps4) | Wild type; marine isolate | This study |
| *Pseudoalteromonas* spp. (Ps6) | Wild type; marine isolate | This study |
| *Pseudoalteromonas* spp. (Ps16) | Wild type; marine isolate | This study |
| *Pseudoalteromonas* spp. (Ps18) | Wild type; marine isolate | This study |
| *Pseudoalteromonas* spp. (Ps19) | Wild type; marine isolate | This study |
| *Pseudoalteromonas gelatinilytica* (Pg24) | Wild type; marine isolate | This study |
| *Pseudomonas zhaodongensis* (Pz15) | Wild type; marine isolate | This study |
| *Pseudomonas* spp. (Ps17) | Wild type; marine isolate | This study |
| *Ruegeria pomeroyi* DSS-3-B (Rp3) | Wild type | M.A. Moran, U Georgia |
| *Vibrio* spp. (Vs5) | Wild type; marine isolate | This study |
| *Vibrio* spp. (Vs9) | Wild type; marine isolate | This study |
| *Vibrio* spp. (Vs8) | Wild type; marine isolate | This study |
| *Vibrio* spp. (Vs13) | Wild type; marine isolate | This study |
| *Vibrio* spp. (Vs14) | Wild type; marine isolate | This study |
| **Plasmids** | | |
| pMRP9 | $Ap^r$, *gfp* | Matthew R. Parsek, U. Washington |

[1] $Ap^r$ = ampicillin resistance

### Supplementary video captions

**Video S1:** Composite video of *Pseudoalteromonas marina (*Pm7*), Pseudoalteromonas marina (*Pm11*)* and *Pseudoalteromonas tetradonis (*Pt12*)* degrading phytoplankton lipids. All three bacteria belong to cluster I (Fig. 2). Note the loss of lipid autofluorescence over the total video duration of 114 h. Scale bar is 100 µm.

https://www.dropbox.com/s/hwx5fxi75js6g00/Supplementary%20Video%201.avi?dl=0

**Video S2:** Composite video of phytoplankton lipid degradation by *Vibrio* spp. (Vs8) and *P. zhaodongensis* (Pz15)*,* two cluster III representatives (Fig. 2). Note the loss of lipid autofluorescence over the total video duration of 114 h. Scale bar is 100 µm.

https://www.dropbox.com/s/ypa7jfp6f5x8mkx/Supplementary%20Video%202.avi?dl=0

**Video S3**: Composite video of phytoplankton lipid degradation by *R. pomeroyi* DSS-3-B (Rp3), *Pseudoalteromonas spp. (*Ps19*)* and *Pseudoalteromonas mariniglutinosa* (Pm22). While Rp3 is a cluster III representative, Ps19 and Pm22 are cluster IV representatives. Note the absence of visible lipid degradation by these three bacteria over the course of 114h of incubation. Scale bar is 100 µm.

https://www.dropbox.com/s/lpxbg3r1fz51um4/Supplementary%20Video%203.avi?dl=0

**Video S4**: **Video sequence of the concentration-dependent quenching of chlorophyll fluorescence within a lipid droplet**. A droplet containing highly concentrated chlorophyll is visible under transmitted visible light, but invisible under epifluorescence ((Ex.: 628/40 nm HBW, Em.: 692/40 nm HBW) because chlorophyll does not emit fluorescent light due to self-quenching. Upon addition of triacylglycerol (trioctanoylglycerol) with a glass pipette, chlorophyll is diluted and becomes unquenched as a result.

https://www.dropbox.com/s/glevtji8epw0l9s/Supplementary%20Video%204.mp4?dl=0

**Video S5**: The chemotactic behavior of *P. zhaodongensis* (Pz15) during phytoplankton lipid degradation. Lipid droplets were video recorded for 10 s every 2 minutes. From each individual video, a maximum intensity projection was created for each time point. The video shows the assembled maximum intensity projections over a period of 48 h. Scale bar is 100 µm.

https://www.dropbox.com/s/sehgsuzvwktnm4a/Supplementary%20Video%204_labelled_scale_bar.avi?dl=0

**Video S6**: The chemotactic behavior of *P. zhaodongensis* (Pz15) during synthetic triacylglycerol degradation. Synthetic TAG droplets (encircled in yellow) were incubated with Pz15::GFP and filmed using fluorescence and phase-contrast microscopy every 30 min for a duration of 60 h. Scale bar is 50 µm.

https://www.dropbox.com/s/aef2azs437ihw3w/Supplementary%20Video%205_labelled_fully.avi?dl=0

**Video S7**: The combined growth of *P. zhaodongensis (*Pz15::GFP*,* green fluorescent) and *A. macleodii* ATTC 27126 (Am2, not fluorescent) in the presence of a phytoplankton lipid droplet. Isolates were imaged in fluorescence (Ex 470 nm / Em 525 nm) and phase-contrast microscopy every 15 min for 47 h. Scale bar is 50 μm.

https://www.dropbox.com/s/fz0b20sb1nt8zn1/Supplementary%20Video%206_labeled.avi?dl=0
